# Continent-wide genomic analysis of the African buffalo (*Syncerus caffer*)

**DOI:** 10.1101/2023.11.12.566748

**Authors:** Andrea Talenti, Toby Wilkinson, Elizabeth A. Cook, Johanneke D. Hemmink, Edith Paxton, Matthew Mutinda, Stephen D. Ngulu, Siddharth Jayaraman, Richard P. Bishop, Isaiah Obara, Thibaut Hourlier, Carlos Garcia Giron, Fergal J. Martin, Michel Labuschagne, Patrick Atimnedi, Anne Nanteza, Julius D. Keyyu, Furaha Mramba, Alexandre Caron, Daniel Cornelis, Philippe Chardonnet, Robert Fyumagwa, Tiziana Lembo, Harriet K. Auty, Johan Michaux, Nathalie Smitz, Philip Toye, Christelle Robert, James G.D. Prendergast, Liam J. Morrison

**Author notes:** Joint first authors (Andrea Talenti, Toby Wilkinson). Joint last authors (Christelle Robert, James G.D. Prendergast, Liam J. Morrison).

## Abstract

The African buffalo (*Syncerus caffer*) is a wild bovid with a historical distribution across much of sub-Saharan Africa. Genomic analysis can provide insights into the evolutionary history of the species, and the key selective pressures shaping populations, including assessment of population level differentiation, population fragmentation, and population genetic structure. In this study we generated the highest quality *de novo* genome assembly (2.65 Gb, scaffold N50 69.17 Mb) of African buffalo to date, and sequenced a further 195 genomes from across the species distribution. Principal component and admixture analyses provided surprisingly little support for the currently described four subspecies, but indicated three main lineages, in Western/Central, Eastern and Southern Africa, respectively. Estimating Effective Migration Surfaces analysis suggested that geographical barriers have played a significant role in shaping gene flow and the population structure. Estimated effective population sizes indicated a substantial drop occurring in all populations 5-10,000 years ago, coinciding with the increase in human populations. Finally, signatures of selection were enriched for key genes associated with the immune response, suggesting infectious disease exert a substantial selective pressure upon the African buffalo. These findings have important implications for understanding bovid evolution, buffalo conservation and population management.

## Introduction

The African buffalo, *Syncerus caffer*, is a key member of the charismatic African megafauna, and was historically distributed across sub-Saharan Africa, inhabiting a diverse range of habitats from dry savannah to montane rainforest. Over the past century the population density and distribution has been much reduced. The population range has also become increasingly fragmented due to man-made pressures, resulting in approximately 70% of the global population being restricted to protected areas [1–3].

The species has been historically divided into varying numbers of subspecies based upon distribution, habitat and morphology, the most recent update of the IUCN Red List recognising *S. caffer caffer* (Eastern and Southern African savannah), *S. c. brachyceros* (Western African savannah), *S. c. aequinoctialis* (Central African savannah), and *S. c. nanus* (Western and Central African forest) [4]. The genetic understanding of population diversity and structure across the species range mostly derives from the application of low resolution tools, such as mitochondrial D-loop sequences, microsatellites and mitogenomes [5–7], with a more recent study using genome-wide single-nucleotide polymorphisms (SNPs) [8]. The two studies to analyse diversity at the genome level, focused on South African *S. c. caffer* animals (n=40) in protected areas [9] and *S. c. caffer* populations (n=59) from East and Southern Africa [10]. These studies have collectively highlighted that the current subspecies classification may not be supported by genetic data, and that there is striking population substructuring within and between the putative subspecies. They have also indicated concerns with respect to low effective population sizes in increasingly isolated populations in some African regions. Improved genetic tools can potentially contribute to conservation management strategies, both in terms of restoring connectivity between relevant populations in order to improve or restore genetic diversity, and avoiding loss of genetic integrity (i.e. maintenance of genetic diversity relevant to local environmental adaptation) through uninformed population mixing (e.g. translocations) [6, 11, 12].

As well as being an iconic species of African wildlife, the African buffalo is the closest bovid relative of domesticated cattle (*Bos taurus taurus & Bos taurus indicus*) in Africa. The African buffalo has co-evolved in Africa with pathogens responsible for important and impactful diseases of cattle such as animal African trypanosomiasis [13] and foot and mouth disease virus (FMD) [14, 15]. For trypanosomiasis, in contrast to the often devastating impact that infection has on cattle, African buffalo are largely tolerant, displaying much less severe clinical signs (e.g. [16, 17]). Additionally, African buffalo are the primary host for the tick-borne protozoan *Theileria parva*, the causative agent of East Coast fever, an often deadly disease in cattle that is asymptomatic in buffalo [18]. These diseases have impeded productivity and the expansion of African pastoralists and their cattle for centuries [19, 20]. During the colonial era, European cattle also brought with them diseases then exotic to Africa, such as rinderpest, brucellosis and bovine tuberculosis [21], to which African buffalo are susceptible. African buffalo and cattle co-exist today across many wildlife/livestock interfaces that enhance mutual pathogen transmission [22], and this can result in imposition of strict veterinary controls at these interfaces that often impact local livelihoods and conservation efforts (e.g., [23, 24]). This makes the buffalo particularly interesting in terms of host-pathogen coevolution and potentially providing a route to identifying host genes and pathways relevant to controlling these diseases in livestock.

This study aimed to develop a reference genome for the African buffalo, as a foundation to analyse the population genomic structure across the current distribution of the species in sub-Saharan Africa. Two reference genomes have previously been published, but were generated via short read sequencing, resulting in relatively fragmented final genome assemblies (scaffold N50s of 2.40 Mb and 2.32 Mb, respectively) [25, 26]. Using a combination of long read (PacBio) and Hi-C sequencing, we generated and *de novo* assembled a substantially higher quality and more contiguous reference genome of 2.65 Gb, with a scaffold N50 of 69.17 Mb. We then sequenced the genomes of 196 African buffalo samples from across the current species distribution, which enabled the analysis of genetic substructure, admixture between populations, and effective population sizes. We also assessed *S. caffer* genomes for signatures of selection, highlighting genes that may be responsible for environmental adaptation, in particular against diseases important for both buffalo and cattle.

## Results

### Assembly statistics

We first generated a *de novo S. c. caffer* reference genome from a male buffalo (OPB4) sampled in Ol Pejeta Conservancy, Kenya, providing the foundation to enable the characterisation of the genetic diversity of African buffalo populations both in terms of their geographic regions and habitats and their current subspecies classification. We applied a deep sequencing strategy, based on a combination of 60× long read (PacBio) and 75× short read (Illumina) reads, to generate a *de novo* reference genome ensuring high per base sequence quality and consensus to achieve good genome contiguity, with an N50 of 69.16Mb. The long reads were assembled using FALCON (Dovetail Genomics) and polished using Arrow. Contigs were then scaffolded using ∼393 million 2×150bp Illumina reads of HiC data, using the HiRise software. Gaps in the draft genome were addressed using PBJelly [27]. Finally, Pilon [28] was used for sequential rounds of polishing, each of which was assessed for its resulting assembly quality over previous rounds. The genome following four rounds of polishing displayed the highest assembly statistics, with a total of 3,351 scaffolds, a total length of 2.65 Gb (comparable to 2.72 Gb for the *Bos taurus* genome), a scaffold N50 of 69.16 Mb and a quality value (QV) of 35.9, indicating ∼1 error every 5,000 bp. The assembly statistics are summarised in Figure 1 and Supplementary Data 1.

**Figure 1.**
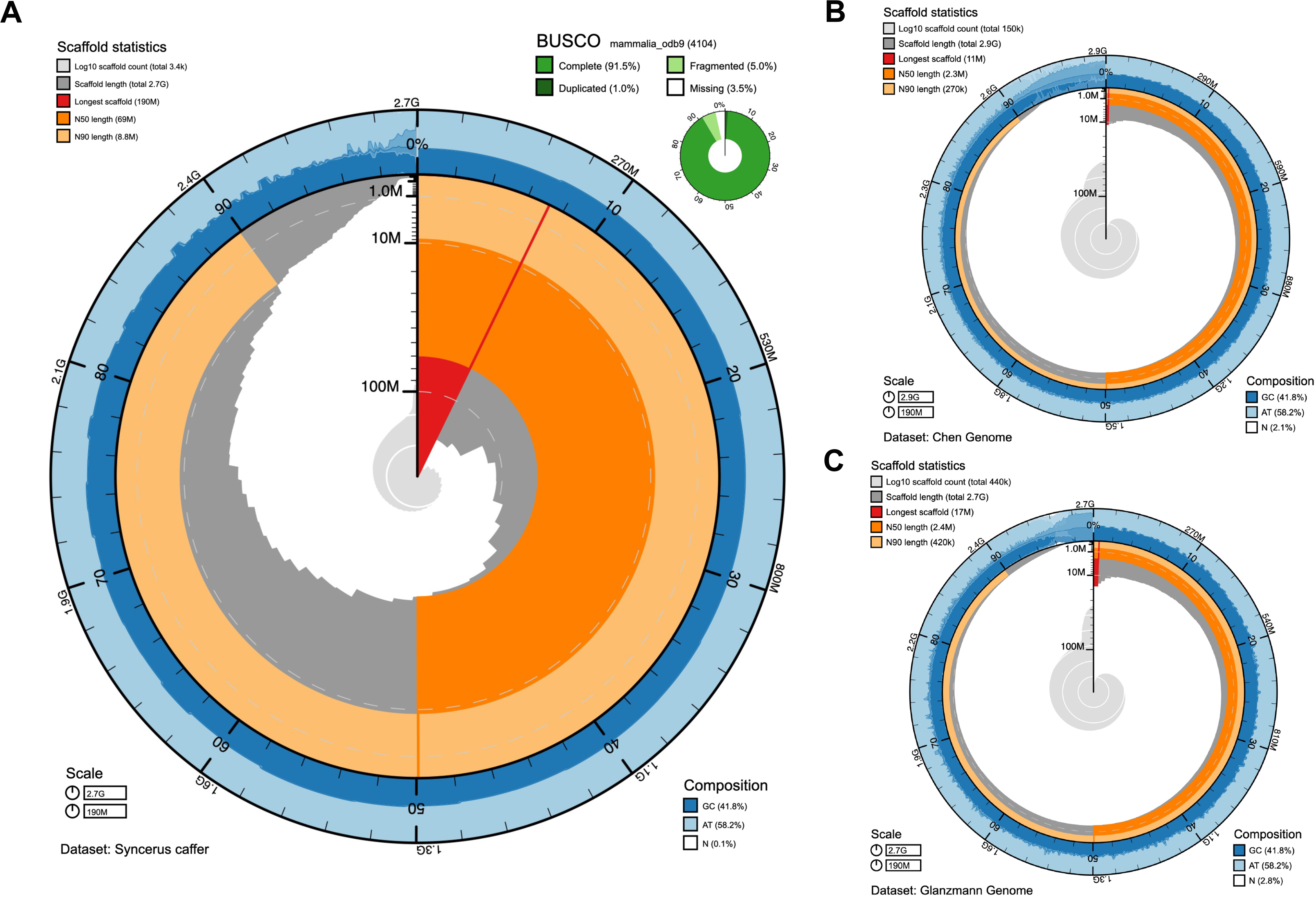
Genome assembly metrics. The BlobToolKit Snailplot shows N50 metrics and BUSCO gene completeness. A) *Syncerus caffer de novo* assembly. The main plot represents the full genome length of 2.65 Gb. The distribution of scaffold length is shown in dark grey with the plot radius scaled to the longest chromosome present in the assembly (190 Mb, shown in red). Dark and light orange sections represent N50 and N90 (69Mb and 8.8Mb), respectively. The pale grey spiral shows the cumulative scaffold count on a log scale with white scale lines showing successive orders of magnitude. The blue/pale-blue/white ring graph shows the distribution of GC, AT and N percentages for the given range in the main plot. A summary of complete, fragmented, duplicated and missing BUSCO genes in the mammalia_odb9 set is shown in the top right. B) Chen et al published genome. BlobToolKit Snailplot representing the *S. caffer* assembly as presented by Chen et al. [26] C) Glanzmann et al published genome. BlobToolKit Snailplot representing the *S. caffer* assembly as presented by Glanzmann et al. [25] The plot radius for both the Chen and Glanzmann genomes has been scaled to the maximum contig length (190Mb) in the *S. caffer* genome assembled here to enable comparison of metrics.

Previous African buffalo reference genomes, generated by Glanzmann et al. [25] and Chen et al. [26], were based solely on Illumina short read sequencing, which led to highly fragmented assemblies of 442,401 scaffolds with a scaffold N50 of 2.40 Mb, and 150,000 scaffolds with an N50 of 2.30 Mb, respectively. These very fragmented assemblies provided limited scope for downstream analysis of variants and their predicted effects on functional regions, i.e. annotated genes and regulatory regions (a comparison of the three genome assemblies is illustrated in Figure 1).

### Transcriptome analyses and genome annotation

To enable in depth characterisation of the African buffalo transcriptome and to facilitate the annotation of gene isoforms, we performed full length isoform sequencing (Iso-Seq) across samples from six different tissues (prescapular lymph node, testis, liver, kidney, lung and spleen) collected from the same animal for which the genome was assembled (OPB4). In total 51,521 distinct, high quality isoforms (defined as being supported by at least two full length reads and with >99% base composition accuracy) were detected across these samples (median of 11,520 per tissue, maximum of 27,271 in the testis). Complementing these data, we also generated Illumina RNA-seq data, from the same animal, from eight tissues (heart, prescapular and inguinal lymph nodes, testis, liver, kidney, lung and spleen). All transcriptomic data were deposited to ENA (https://www.ebi.ac.uk/ena/browser/view/GCA_902825105.1) with accession numbers PRJEB36587 and PRJEB36588 for RNA-seq and Iso-Seq, respectively. Together these data have been used to provide a high quality annotation of the buffalo assembly which can be accessed through the Ensembl Rapid Release genome browser: https://rapid.ensembl.org/Syncerus_caffer_GCA_902825105.1.

### African buffalo-specific sequence

After aligning the African buffalo genome to eight high quality assemblies of four different Bovidae species (cattle, water buffalo, yak and goat [29–34]), portions of the *S. caffer* genome that did not match any regions in the other assemblies were ascertained. This process identified a total of 24,336,918 intervals, for a total of 145,050,830 bp of sequence not identified in the other eight assemblies. This includes both small variations (e.g. SNPs, small indels), unplaced contigs without alignments to any other genome, and large portions of the genome lacking any alignment.

We then refined the region selection by filtering out shorter intervals (<60 bp) and regions defined as too close to a telomere (<10 Kb) or to a gap (<1 Kb), leaving a total of 113,654,400 bp in 81,357 fragments longer than 60 bp, which were neither telomeric nor neighbouring an assembly gap. These regions have an average length of 1,397bp (3772.4 bp SD) and a median size of 286bp (min. 61bp, max 308,890bp). The majority of the regions (74,659 fragments accounting for 112,762,919 bp) represent sequence not found in any of the other species genomes considered in the study, whereas the remaining are classified as divergent haplotypes. Of the 113Mb, a total of 64.9Mb (57.1%) are putatively identified as repeats using RED [35]. To rule out the possibility of these novel regions being due to contamination, we confirmed the coverage of these regions was consistent with the rest of the genome, using short-read whole genome sequencing data from 46 samples from the population analysis (see section below; Supplementary Figure 1).

HOMER analysis considered 4,286/7,096 sequences with less than 60% of masked nucleotides. These sequences presented 38 motif types enriched (P-value <1e-5), such as the FOSL2/MA0478.1/Jaspar (0.661) motif, a negative regulatory sequence in the differentiation-sensitive adipocyte gene (aP2), a motif identified as a transcriptional enhancer for the Gibbon ape leukaemia virus, and which is also in a region of the human immunodeficiency virus (HIV) [36]. We performed the feature analysis on the annotation generated by Ensembl from the Iso-Seq sequencing data previously described. We identified 7,096 annotated genes and 131 pseudogenes overlapping the novel regions, of which 583 genes, 194 ncRNA genes and 71 pseudogenes were entirely included in the identified regions (Supplementary Table 1). A total of 317 of 583 genes had at least one biological term annotated. GO terms definitions were fetched using the goatools python package [37]. Out of 4,088 terms in the background dataset, 17 (15 GO terms and 2 KEGG pathways) were found significantly enriched. Among the significant terms was the defence response GO term (GO:0006952, FDR-corrected P-value: 0.0189, Supplementary Table 1), described as the response triggered by the presence of a foreign body.

## Population genetics

To better understand African buffalo genetic diversity, we generated short read sequencing data for a further 195 animals deriving from across the continental range of the species (at a coverage of 15× for 146 samples, and 30× for 50 samples; Table 1 & Figure 2A; for full sample list and metadata see Supplementary Table 2). This included samples from the currently described four subspecies; *S. c. caffer*, *S. c. nanus*, *S. c. brachyceros* and *S. c. aequinoctialis* (Table 1 & Figure 2A), and two putative *S. c. nanus* and *S. c. aequinoctialis* hybrids (based upon morphology and geography at time of sampling – labelled as ‘intermediates’). Together, these samples derived from 21 sites/localities or protected areas across 12 different countries. We performed phylogenetic and population analyses including only samples with a high call rate (>85%), and analysing only the biallelic polymorphic SNPs (minor allele frequency >5%), as well as only considering unrelated individuals (samples fourth degree or greater).

**Table 1.**
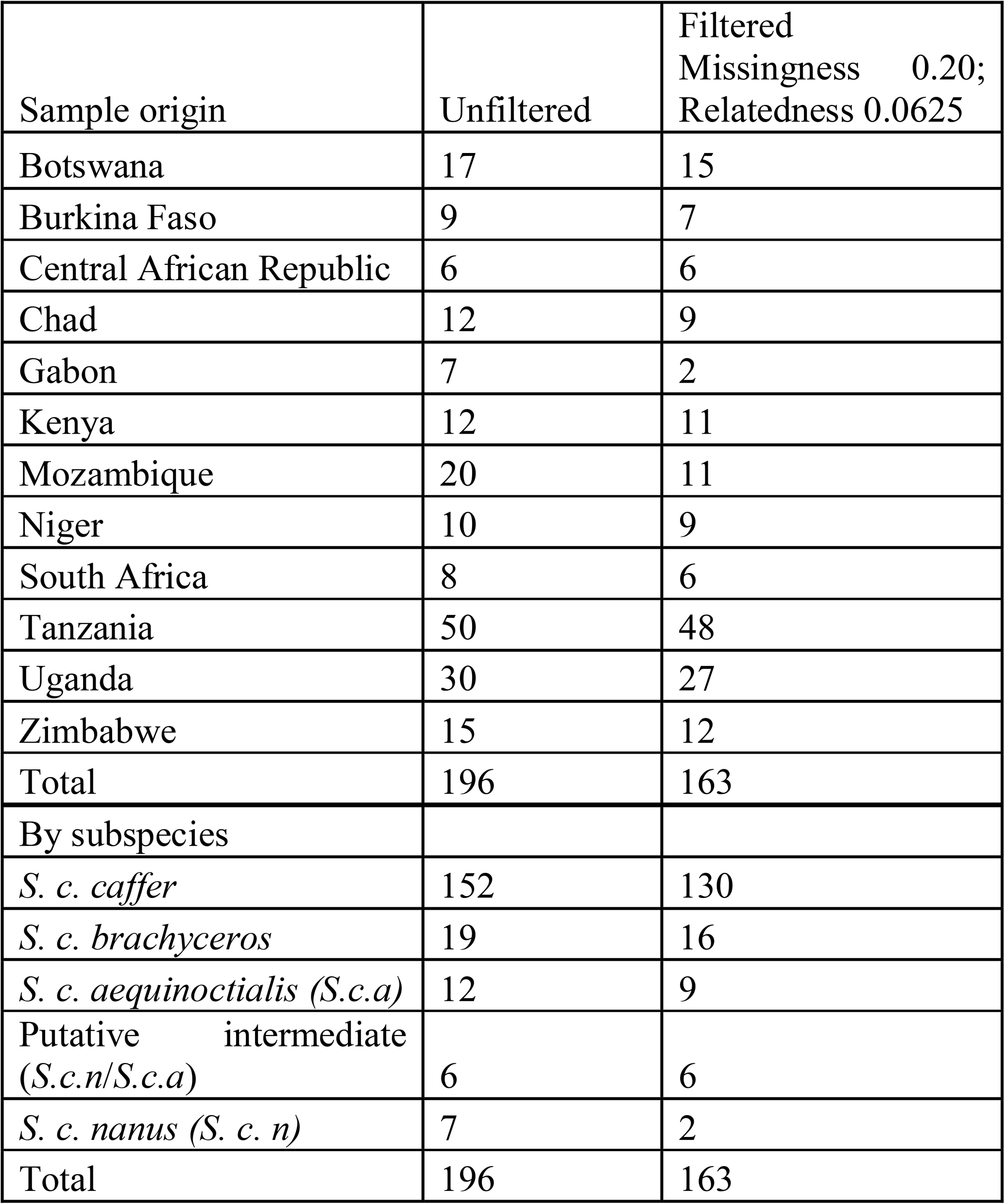
Sample number by country, subspecies and pre– and post-data filtering.

**Figure 2.**
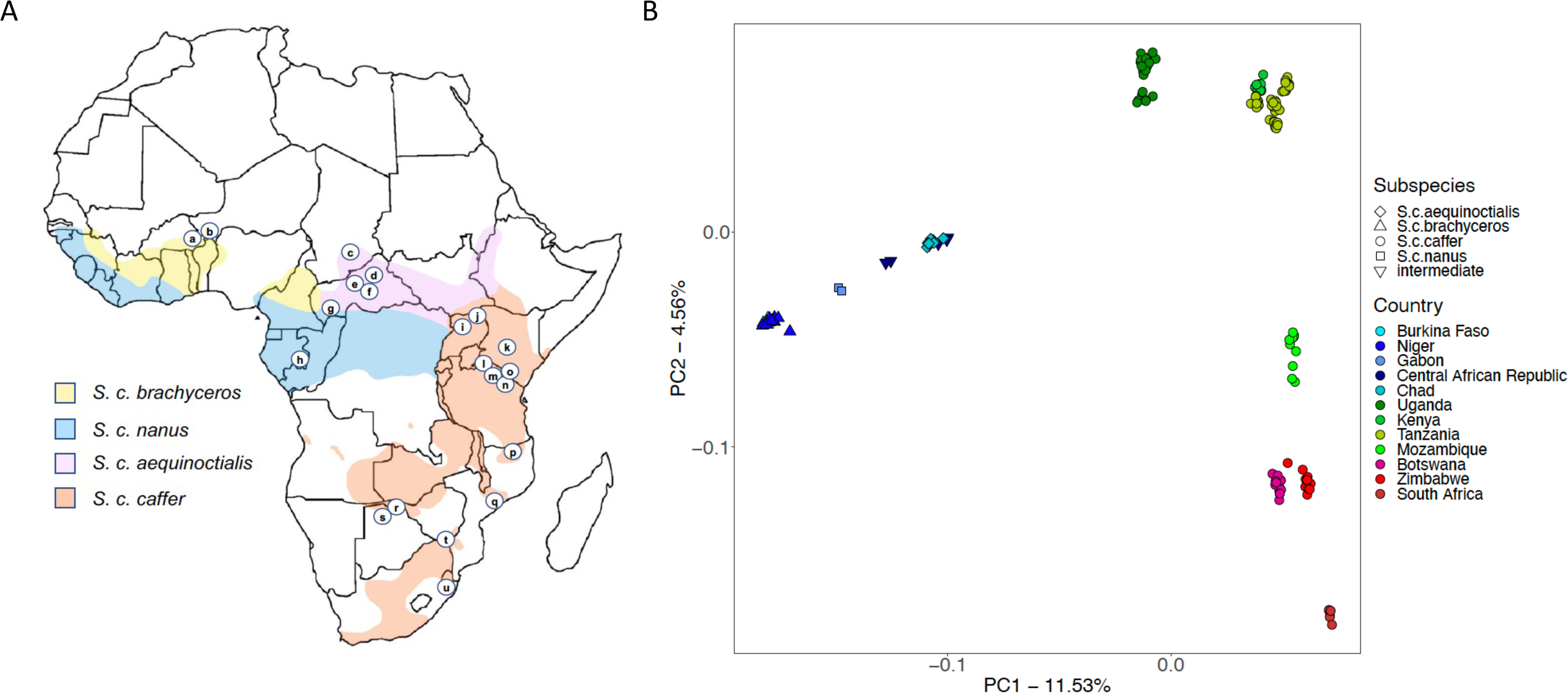
(A) The sampling locations of African buffalo samples sequenced in the current study (circled letters), mapped on to the approximate current distribution of the four subspecies. a: Singou and Pama Game Reserves (GR)/Arli National Park (NP) complex, Burkina Faso (n=10 samples [before data filtering]); b: W NP, Niger (n=10); c: Zakouma NP, Chad (n=13), d: Manovo-Gounda-St. Floris NP, Central African Republic (CAR; n=2); e: Bamingui-Bangoran NP, CAR (n=2); f: Sangba, CAR (n=1); g: N’Gotto Forest Reserve, CAR (n=2); h: Lekedi NP, Gabon (n=8); i: Murchison Falls NP, Uganda (n=13), j: Kidepo NP, Uganda (n=20); k: Ol Pejeta Game Reserve, Kenya (n=12); l: Serengeti NP, Tanzania (n=15); m: Ngorongoro Conservation Area, Tanzania (n=15); n: Tarangire NP, Tanzania (n=10); o: Arusha NP, Tanzania (n=10); p: Niassa National Reserve (NR), Mozambique (n=9); q: Marromeu NR, Mozambique (n=9); r: Chobe NP, Botswana (n=9); s: Okavango Delta, Botswana (n=9); t: Gonarezhou NP/Crook’s Corner, Zimbabwe (n=18), u: Hluhluwe-Umfolozi NP, South Africa (n=8; for full sample data see Supplementary Table 2). (B) Principal Component Analysis of population samples, with data for components 1 and 2 illustrated. Samples are coloured by country of origin, with different symbols indicating the previously recognised subspecies.

As can be seen in Figure 2B the genetic relationships between the samples largely mirrors their geographic origin, with the first principal component (PC1) reflecting differentiation between samples from Eastern/Southern and Western Africa, which corresponds to a split between the Western/Central African subspecies (*S. c. aequinoctialis*, *S. c. brachyceros* and *S. c. nanus*) and Eastern/Southern African *S. c. caffer.* The second component (PC2) correlates with differentiation between *S. c. caffer* samples from the Northern part of the subspecies’ range (Kenya, Tanzania, Uganda) compared to *S. c. caffer* samples from Southern Africa. Notably, there was a clear signature of geography within the *S. c. caffer* data, with each geographic sub-population forming a distinct cluster in the PCA and a cline observed from Uganda to Kenya and Tanzania in the North, through Mozambique to samples from Botswana and Zimbabwe, and finally South Africa in the South. In Western/Central Africa, *S. c. aequinoctialis*, *S. c. brachyceros* and *S. c. nanus* sub-populations also formed separate clusters, although the *S. c. nanus* and *S. c. brachyceros* populations clustered closely together. Samples were initially grouped by sub-species and country of sampling. However, based on PCA results, the Tanzania and Kenya, and Botswana and Zimbabwe samples were grouped together, reflecting their geographic proximity. This resulted in nine subgroups for downstream analyses; referred to hereafter as *S. c brachyceros*, *S. c. nanus*, *S. c. aequinoctialis*, intermediate (putative hybrids between *S. c. nanus*, *S. c. aequinoctialis*), *S. c. caffer* Uganda, *S. c. caffe*r Kenya/Tanzania, *S. c. caffer* Mozambique, *S. c. caffer* Zimbabwe/Botswana and *S. c. caffer* South Africa. Population sample sizes post-filtering ranged from 2 for the *S. c. nanus* spp to 48 for the *S. c. caffer* from Tanzania (see Table 1), leaving a total of 163 samples for the phylogenetic analyses.

In order to explore the relationship between these populations further, and to mitigate the different sample size between subpopulations resulting in over-representation of population-specific variation in the dataset as far as possible [38], we downsampled the larger groups to 15 representative samples (for those with less, all samples were included). Since the *S. c. brachyceros* population had a total of 16 samples, we did not perform any reduction on this population. This resulted in a subset of 95 individuals to be considered for the population genetic analyses (see Supplementary Table 2 for samples included in these analyses). As shown in the principal components analysis (PCA) pre– and post-reduction (Supplementary Figure 2), the general structure of the sample was not affected by the subsampling.

Bootstrapped admixture analyses identified three clusters as a parsimonious solution for the number of subpopulations (i.e. lowest iteration number and reduced increase in CV error; Supplementary Figure 3), representing the East, West and Southern African high level groupings. At K=9 the clusters recapitulate the nine sub-groupings described above (see Supplementary Figure 4 for admixture results at multiple K). The same high-level structure is reflected in the 100-bootstrap identity-by-state phylogenetic tree (Figure 3B).

**Figure 3.**
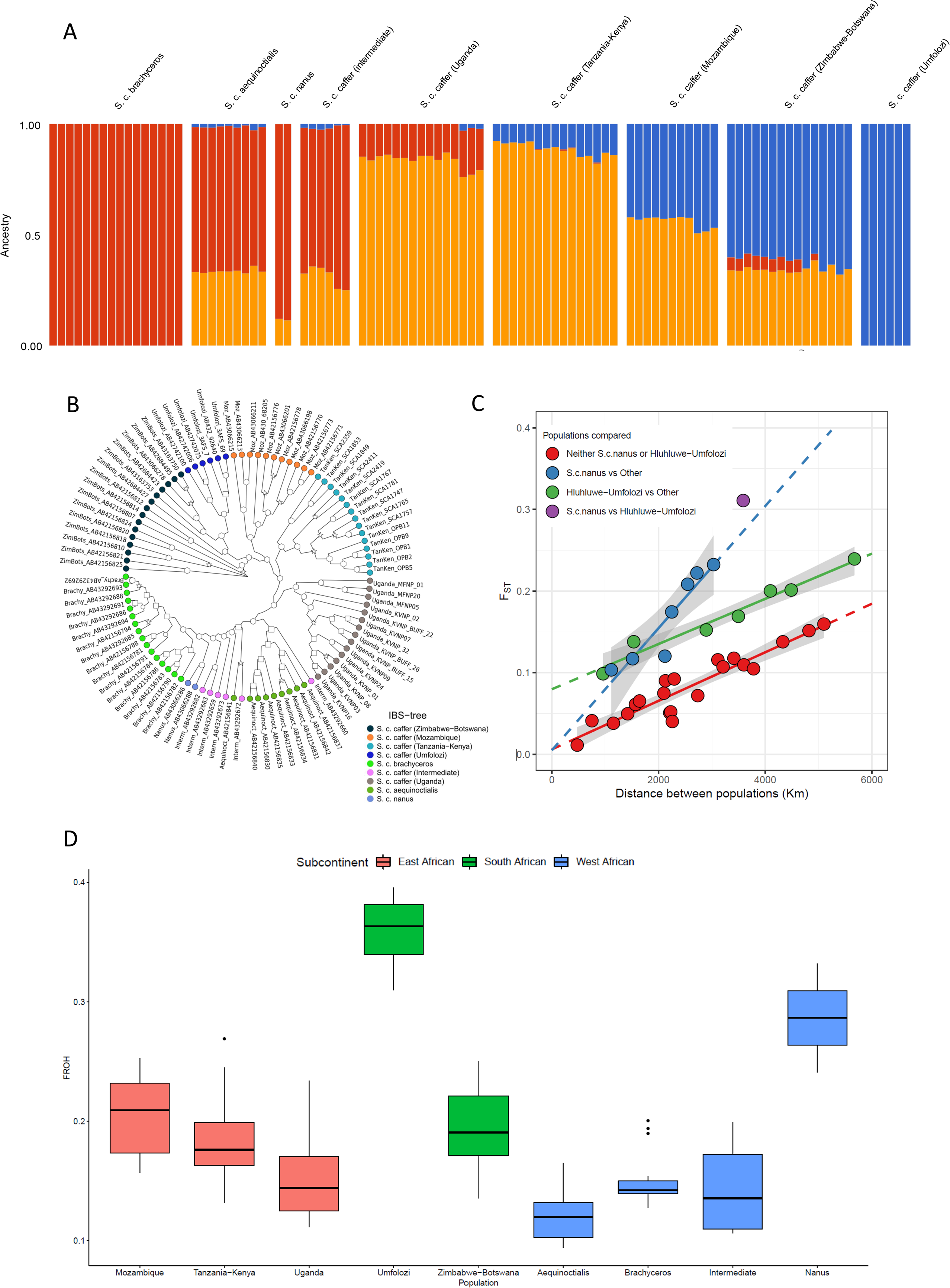
Population genetic analyses based on genome sequences: A) Admixture analysis for K = 3 showing the three major clusters of diversity (Western/Central Africa in red, Eastern Africa in yellow and Southern Africa in blue). B) Neighbour-joining phylogenetic tree from the Identity-by-State (IBS) distances showing the gradient from Southern-to Eastern-to Western/Central-African populations (clockwise from the root). Symbols indicate confidence value for each node (circle is less than 50%; square between 50% and 74%; star between 75% and 89%; hexagon above 90%). C) Isolation by distance (IBD) analysis of African buffalo populations. The F_ST_ values were calculated between all pairs of populations and plotted against their geographic distance apart. Pairwise comparisons involving *S. c. nanus* are indicated in blue, pairwise comparisons involving the Hluhluwe-Umfolozi population are shown in green, the single pairwise comparison comparing *S. c. nanus* and Hluhluwe-Umfolozi in purple, and all remaining pairwise comparisons in red. The predicted pairwise F_ST_ values outside of the observed distances are indicated by dashed lines. D) The proportion of homozygous segments per sample (FROH) indicating that the Hluhluwe-Umfolozi population has unusually high levels of homozygosity.

Comparison of the genetic diversity between all pairs of populations (as represented by the F_ST_ statistic) highlights that this is largely a function of physical distance, i.e. the diversity observed between two populations increases broadly linearly with increasing distance between them (Figure 3C, Mantel test r: 0.65, p=0.0018; underlying F_ST_ data detailed in Supplementary Table 3). However, sub-structure in this isolation-by-distance analysis is observed. After excluding the *S. c. caffer* Hluhluwe-Umfolozi and *S. c. nanus* populations, the relationship is even stronger, and variation in the F_ST_ values between the remaining groups can potentially largely all be explained by the distances between them (red line in Figure 3C, Mantel test r: 0.96, p=0.0013). This is consistent with the idea that these African buffalo have historically formed large continuous groups of populations with differentiation between populations simply reflecting the reduced mating probability with increasing distance. *S. c. nanus*, the forest buffalo, shows an unusually steep increase in differentiation relative to other populations (blue line in Figure 3C). This could be for a variety of reasons, including geographical barriers reducing the gene flow between this group and the others analysed. Animals found at the same location should exhibit little differentiation, and consistent with this, the intercept of the slopes is not significantly different from 0 in these comparisons (both linear regression intercept P>0.4), i.e. when comparing *S. c. nanus* to other populations or the non-*S. c. nanus* and non-*S. c. caffer* Hluhluwe-Umfolozi populations to each other. However, this is not the case for comparisons involving the South African *S. c. caffer* Hluhluwe-Umfolozi population. Under the assumption of a simple linear relationship between genetic differentiation and geographic distance, the predicted level of diversity at a distance of 0 km is significantly higher than 0 (green line in Figure 3C, linear regression intercept P=2.7×10^-4^). This suggests, that unlike in the other population comparisons, there is elevated differentiation between this population and others, above and beyond that expected from their geographic distance apart. This may reflect an isolation event with respect to the Hluhluwe-Umfolozi population.

EEMS analysis (Figure 4A) adds to this picture of continental gene flow, with the Congo river basin likely representing a significant barrier of migration, particularly between Western/Central African *S. c. nanus* and *S. c. caffer* populations in Eastern Africa. The data also suggest that the Rift Valley potentially presents a geographical barrier to gene flow within the African buffalo.

**Figure 4.**
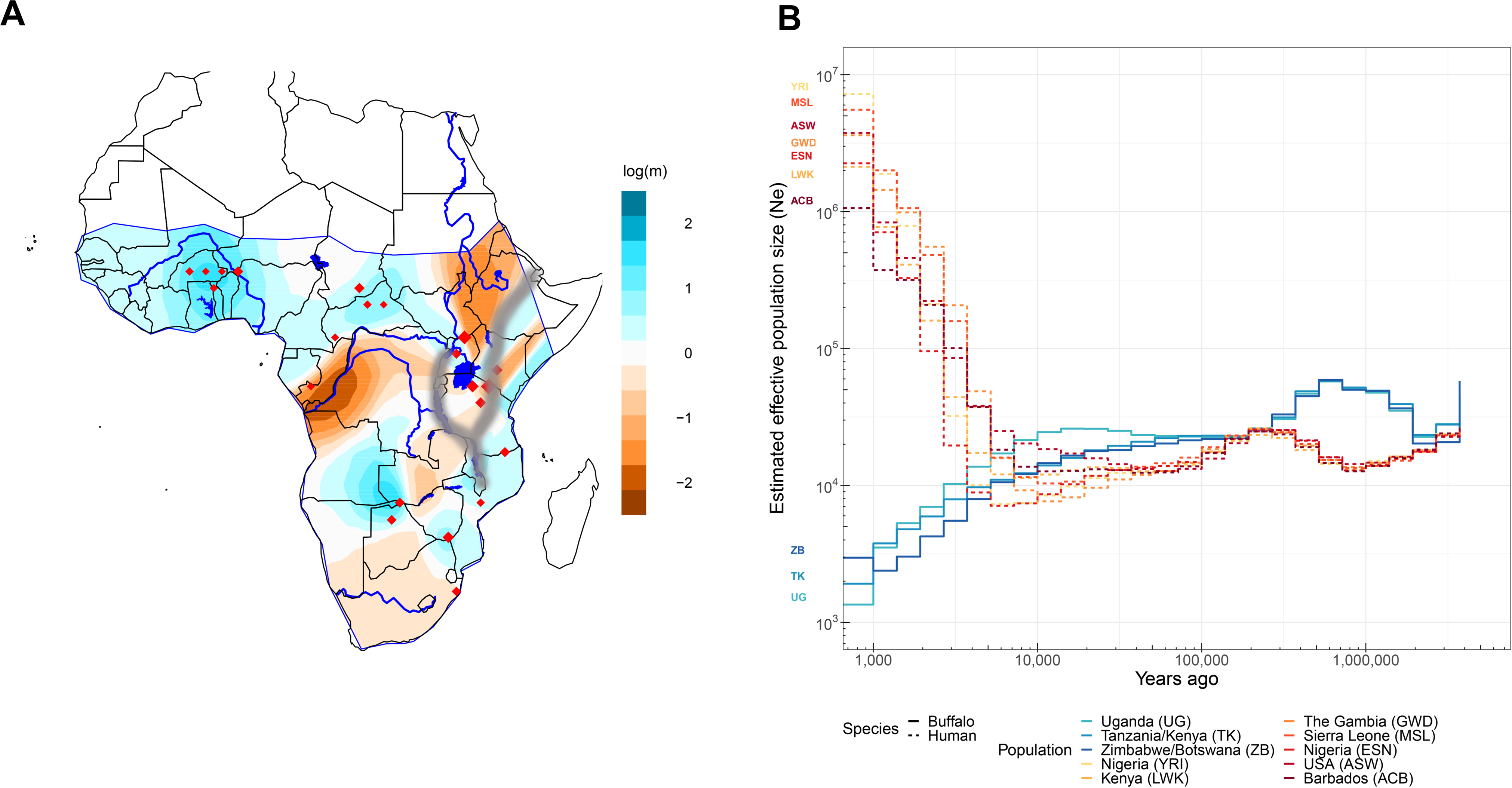
(A) The contour map shows the mean of two independent Estimating Effective Migration Surfaces (EEMS) posterior migration rate estimates between 400 demes modelled over the land surface of Sub-Saharan Africa. A value of 1 (blue) indicates a tenfold greater migration rate over the average; −1 (orange) indicates tenfold lower migration than average. The courses of the major river systems (Niger, Congo, Nile and Orange rivers), as well as water bodies with a surface area greater than 5,000 km^2^ are included to highlight their potential relationships with migratory routes and barriers. Red diamonds indicate geographical location of samples in the dataset. (B) Estimated effective population sizes of African buffalo (solid lines) and human (dashed lines) populations over time. The countries of sampling for each population are indicated in the legend along with the three letter 1,000 Genomes consortium population code for the human data. Only human populations from the 1,000 Genomes consortium dataset of recent African origin are shown.

The Relate software and genome-wide genealogies were used to estimate population-specific population sizes over time for the largest buffalo groupings (*S. c. caffer* only: Uganda, Tanzania/Kenya and Zimbabwe/Botswana – grouped as defined by PCA, admixture and phylogenetic analyses; Figures 2 and 3). As shown in Figure 4 there has been a sharp reduction in the estimated effective population sizes across these groups in the last approximately 10,000 years, broadly mirroring the expansion of human effective population sizes over a similar time-period (Figure 4B). There were not sufficient numbers in all individual populations for robust N_e_ analyses, but for the populations that did have sufficient numbers, contemporary N_e_ estimates were ∼1,300, 2,000 and 3,000 for Uganda, Tanzania/Kenya and Zimbabwe/Botswana, respectively. These data suggest that the effective population sizes of these Eastern and Southern African *S. c. caffer* are above the levels of conservation concern. Coelescence estimates are shown in Supplementary Figure 5.

However, analysis of all populations highlights that the *S. c. nanus* and South African *S. c. caffer* Hluhluwe-Umfolozi samples have high levels of homozygosity (F_ROH_ of 0.29 and 0.36 compared to a range of 0.12 to 0.21 for the other populations; Figure 3D). This is consistent with the known extreme bottlenecks experience by the Hluhluwe-Umfolozi buffalo population [9]; the *S. c. nanus* samples derive from Lekedi NP in Gabon, and we are unaware of historical population-level data that would inform of bottlenecks – while the homozygosity analysis is obviously on individual genomes, with this population we would caution overinterpretation as we only have data from two individuals.

### Selective sweeps

African buffalo are exposed to a range of different environmental pressures across their distributional range, including a range of pathogens that also impact domesticated bovids such as cattle. To investigate selective sweeps between and within the nine population groupings we calculated the XP-EHH and P_R_ Relate Selection Test statistics [39, 40]. Due to being more susceptible to artefactual results deriving from smaller sample sizes than the XP-EHH statistic, the calculation of the P_R_ statistic was restricted to just the populations with more than 20 samples after filtering for relatedness (i.e. the Uganda, Zimbabwe/Botswana and Tanzanian/Kenyan populations). These two tests are complementary in that whereas the XP-EHH statistic tests for differences in haplotype homozygosity between populations, P_R_ characterises the speed of spread of particular genomic lineages within a population, relative to others. Supplementary Table 4 summarises the results of these two tests. In total, 73 loci of elevated XP-EHH levels overlapping a gene were identified in at least one population comparison, and 34 P_R_ significant loci were detected in one of the three studied populations. Of the XP-EHH loci, 9 also overlapped a significant P_R_ peak (Supplementary Table 4). These 9 loci spanned 11 genes, with several having strong links to immune response, including putative killer cell immunoglobulin-like receptor like protein KIR3DP1 (LOC102402296), T cell receptor beta variable 5-1-like (LOC112577699), the major histocompatibility complex gene TRIM26 and N-acetylneuraminic acid phosphatase (NANP). The latter is involved in sialic acid synthesis which in turn is linked to immune response modulation, and NANP has also been observed to be under recent positive selection in both humans and cattle [41, 42]. Two further of these nine genes linked to both XP-EHH and P_R_ peaks in African buffalo were also previously linked to recent positive selection in water buffalo [42], namely myeloid-associated differentiation marker-like (LOC102403696) and tyrosine-protein phosphatase non-receptor type substrate 1-like (SIRPA-like) gene (LOC102396916). LOC102396916 was associated with significant P_R_ peaks in both the Uganda and Tanzania/Kenyan populations and also elevated XP-EHH scores in the South African *S. c. caffer* vs intermediate and *S. c aequinoctialis* populations (Figure 5; Supplementary Figure 6). SIRPA is an immunoglobulin-like cell surface receptor for CD47 (a cell surface protein that is involved in the promotion/regulation of cellular proliferation) and has been associated with a range of infectious diseases, including *Theileria annulata* infection in cattle [43] (*T. annulata* being the causative agent of tropical theileriosis across North Africa and Asia, and is closely related to *Theileria parva* found in Eastern Africa). This gene has previously been identified to be associated with selective sweeps between water buffalo breeds (elevated XP-CLR statistics between Mediterranean and Jaffrabadi, and Pandharpuri and Banni water buffalo breeds [42]). Characterisation of this gene’s expression profile in the water buffalo expression atlas highlighted that it falls within a macrophage-specific cluster of genes [44]. Together these results therefore point towards this gene being a potentially important target of selection across bovids due to its role in immune response. Consequently, five of these nine genes under putative selection in African buffalo show strong links to immune response, with two of the remaining genes being uncharacterised and their function being unknown.

**Figure 5.**
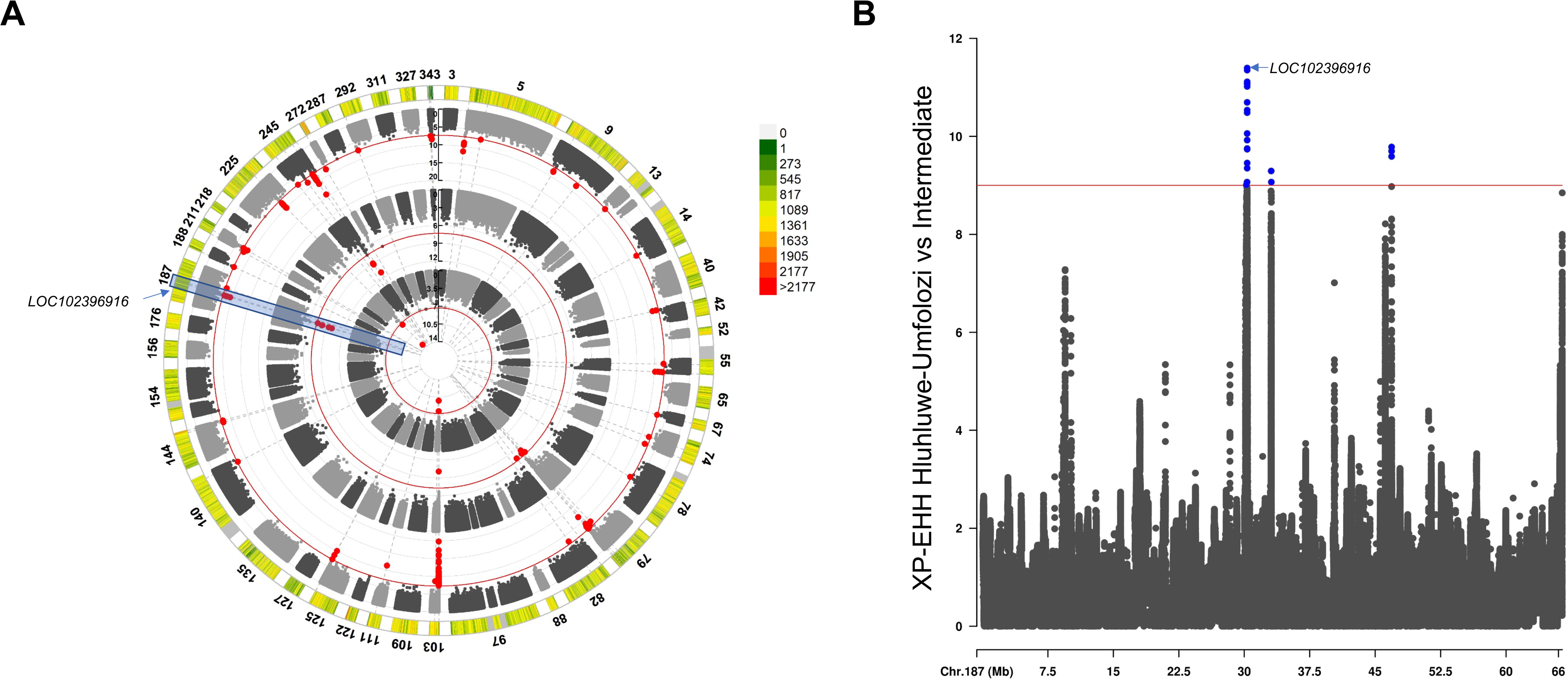
(A) The coloured outermost track and legend indicates the SNP density across 41 large contigs. The next three tracks show the P_R_ scores in the Uganda (centremost), Zimbabwe/Botswana (middle) and Tanzania/Kenya (outer) populations. Red points indicate SNPs with a P-value smaller than 5×10^-8^. The peaks at LOC102396916 are highlighted (B) Absolute XP-EHH scores across Contig 187 for the Hluhluwe-Umfolozi versus intermediate populations indicating the peak also detected at the LOC102396916 locus.

## Discussion

### African buffalo genome

The genome generated in this study represents a substantial improvement on current genomic resources available for *S. caffer*, with greater contiguity and much improved assembly and annotation – this, and the allied gene expression datasets, will hopefully serve as useful resources for the bovid and African buffalo research communities. The genome assembly is currently at the scaffold rather than chromosomal level, and so karyotype and features such as centromeres remain undefined, and the genome also contains Y chromosome and mitochondrial sequences that have not been completely resolved. There is therefore clearly scope for further improvement of the reference genome. An interesting finding was the African buffalo-specific sequence, which was identified after aligning the African buffalo genome to eight existing high quality bovid genome assemblies (cattle, water buffalo, yak and goat [29–34]). *S. caffer* sequences that that did not match any regions in the other assemblies were defined as African buffalo-specific sequence. These sequences were validated by assessing coverage of these African buffalo-specific sequences in randomly selected short read data from the population data, based on the expectation that if these were genuine African buffalo-specific sequence there would be coverage detected in multiple samples, and this was indeed the case. While 57.1% of these African buffalo-specific sequences are repeats, there are 583 genes, 71 pseudogenes, and 194 ncRNAs that are entirely within the identified regions. These were enriched for genes associated with the host defence, and the genes within these regions would clearly be of interest in further studies to identify traits that may be relevant to these African buffalo-specific sequences.

### Population genomic structure: taxonomic insights

The admixture analysis suggests that the *S. caffer* population splits into three high-level lineages, with a further nine subgroupings apparent that correlate with geographical location. There is little support in our data for the current classification of the four IUCN recognised subspecies; *S. c. caffer* (Eastern and Southern African savannah), *S. c. brachyceros* (Western African savannah), *S. c. aequinoctialis* (Central African savannah) and *S. c. nanus* (Western and Central African forest), with *S. c. brachyceros* and *S. c. aequinoctialis* sometimes being lumped and treated as a single subspecies, viz. *S. c. brachyceros*. Historically these classifications have been based on a combination of geographical distribution, habitat preferences and morphological features. *Syncerus c. nanus*, the forest buffalo, is the most divergent morphologically, being on average much smaller, predominantly rufous in colour as opposed to black, and with a different horn shape. From this perspective it is perhaps surprising that we could not detect substantial genetic divergence from the Western/Central African savannah buffalo. However, this finding agrees with previous genetic analyses using mitochondrial D-loop sequence markers, which similarly indicated a lack of support for differentiation between Western/Central African ‘subspecies’ [5]. However, the limited number of samples assigned to *S. c. nanus* did not enable balanced analyses, with in general a smaller number of populations sampled for the Western and Central African regions compared to Eastern and Southern Africa. This may have resulted in some bias in our population analyses. However, we did attempt to mitigate this bias to some extent by reducing populations to 10 samples per population where relevant and possible. Notwithstanding this fact, the present database provides genome-level and –wide resolution on variation (based upon 23,454,419 identified variants relative to the assembled reference genome); a much more robust basis for identifying genetic differentiation than previous methods used to identify genetic substructuring in this species. These insights have parallels with previous genome/multilocus genetic data studies on African ungulates with similar pan-Sub-Saharan distribution, the giraffe (*Giraffa camelopardis*) and zebra (*Equus quagga*), which indicated a lack of correlation of genetic data with morphology-based speciation, in those cases resulting in the identification of cryptic speciation [45, 46].

Admixture and EEMS analyses indicate that the population genomic structure is shaped by geographical barriers, which limit where migration and therefore where lineage and population mixing can happen. This is evidenced by Ugandan buffalo demonstrating ancestry from both Eastern and Western African populations, and there being some signal of East African ancestry in Central African buffalo (*S. c. aequinoctialis*, *S. c. nanus* and intermediate; Figure 3A and Supplementary Figure 4). Both admixture and EEMS data indicate that Uganda is likely to act as an interface zone between these lineages, although further sampling in relevant populations (for example, known buffalo populations in Eastern CAR and DRC, South Sudan and Western Ethiopia) would help resolve the extent of genetic flow. EEMS analyses suggests that the divergence between the East and West lineages was most likely driven by geography, with the Congo Basin and River effectively creating a barrier to North-South gene flow in the West of the continent, and Uganda being the pinch point at which Central African savannah and forest populations can intersect with Eastern African savannah buffalo.

The driving forces shaping the differentiation between Northern and Southern populations of *S. c. caffer* (i.e. between the Kenyan, Ugandan and Tanzanian cluster and the Mozambique, Botswana, Zimbabwe and South Africa cluster) is less clear from our analyses. A potential role of the Great Rift Valley acting as historical barrier to gene flow has been suggested within other large savannah mammals [47–49]. However, all Tanzanian samples included in the present study originated from the North of the country (the closest population in the Southern cluster being Niassa Special Reserve in Mozambique –approximately 1,000 km from the Northern Tanzanian parks); additional samples from Central and Southern Tanzania where substantial buffalo populations exist (e.g. in Ruaha and Nyerere NPs) could potentially identify animals that are genetically intermediate between the ‘Northern’ and ‘Southern’ clusters, and reveal that there is a steady cline of differentiation within *S. c. caffer* from North to South, as supported by the isolation-by-distance analysis. The data are consistent with the findings of a previous genomic study of *S. c. caffer* across its range, which also concluded that there was a primary split between northern and southern *S. c. caffer* populations approximately 50,000 years ago, followed by gene flow [10].

### Effective population sizes

Although effective population size estimates are difficult to estimate accurately and can be confounded by population structure, the effective population size data interestingly suggests a coincident drop in N_e_ with the rise in human N_e_ (obtained through the 1,000 Genomes data [50]). This is observed in similar analyses applied to both other individual African ungulates (giraffe) [51] and collated global ruminant data [26]. In the case of African buffalo, previous studies based on both microsatellite and mitochondrial DNA data have suggested an expansion approximately 80,000 years ago coincident with the spread of grassland habitat, which was followed by a significant decline ∼3-7,000 years ago, probably resulting from an overall increase in arid areas across Africa that are inhospitable to African buffalo [7, 52, 53] – our findings are also consistent with these conclusions. For the African buffalo, it was anticipated that the greater resolution provided by genomic data may detect a drop in N_e_ observed as a result of the rinderpest virus epidemic of the 1890s [54], which anecdotally caused very high mortality of the buffalo populations through Eastern and Southern Africa in particular [55, 56]. However, given the relatively recent timing of the rinderpest epidemic and the fact that the N_e_ was reducing across the relevant timeframe in our analysis, from the genome data we are not able to infer the impact of rinderpest upon population sizes. Other analyses using lower resolution genetic markers [53, 57, 58] were also not able to detect a drop in N_e_ that correlated with the timing of the rinderpest epidemic, although a recent genomic study using samples from *S. c caffer* did identify a very significant drop in N_e_ over the past 500 years, which could plausibly be explained by rinderpest [10] – notably the decline was particularly steep in samples from Hluhluwe-Umfolozi. While we did not have sufficient numbers in each population to robustly test N_e_, for the closely related groupings of Uganda, Tanzania/Kenya and Zimbabwe/Botswana N_e_ estimates were approximately of 1300, 2,000 and 3,000 individuals, respectively. In these clusters at least, there is limited evidence for inbreeding depression, in agreement with previous studies [6]. However, the *S. c. nanus* and South African *S. c. caffer* Hluhluwe-Umfolozi samples showed high levels of homozygosity, meaning that further population-specific work is required in order to assess inbreeding risk. The *S. c. caffer* Hluhluwe-Umfolozi population is known to derive from very small number of founder animals, and our finding is in agreement with previous data that has indicated high inbreeding coefficients and low genome-wide heterozygosity levels in this population [9, 10].

While we have very limited numbers of *S. c. nanus* samples, the finding of high levels of homozygosity may be explained by the very different features of forest buffalo behaviour, in that relative to savannah buffalo forest buffalo have smaller home ranges, shorter daily movements, negligible seasonal movements and live in significantly smaller group sizes [2]. This is linked to the forest habitat likely generally acting as a greater barrier to gene flow than savannah environments, limiting migration/dispersal and resulting in comparatively small and isolated populations [5]. Genetic diversity metrics such as heterozygosity/homozygosity and effective population size will clearly be an important feature for future studies, particularly where there are increasingly fragmented and isolated populations, as is the case for the West African Savannah buffalo.

### Selective Sweeps

The selective sweep analyses identified tyrosine-protein phosphatase non-receptor type substrate 1-like (SIRPA-like) as being under selection, independently detected using two distinct and complementary methodologies (P_R_ and XP-EHH), and across several population groupings (Ugandan, Tanzanian/Kenyan, South African *S. c. caffer*, intermediate and *S. c aequinoctialis* populations). The same locus was identified in selective sweep analyses of the Asian buffalo *Bubalus bubalis* [42], and expression analysis in this species identified upregulated gene expression in a macrophage-specific cluster. Interestingly SIRPA has been associated with *Theileria annulata* infection in cattle [43], and its gene expression has been shown in independent studies to be significantly upregulated in host cells following infection and the cellular transformation associated with *T. annulata* infection [59, 60]. While SIRPA will clearly be involved in the immune response to other pathogens, it is notable that *B. bubalis* is the primary host of *T. annulata* (the tick-borne causative agent of tropical theileriosis across North Africa and Asia). *Syncerus caffer* is similarly the primary host for the related parasite *Theileria parva* (and the related *Theileria* sp. buffalo [61]), and it is therefore plausible to link the described function of this gene with the long co-existence and co-evolution of *S. caffer* with *T. parva.* Although only the Ugandan, Tanzanian/Kenyan and South African *S. c. caffer* populations are within the current distribution of the tick vector (*Rhipicephalus appendiculatus*) of *T. parva*, the historical range and selection of *T. parva* cannot likely be inferred by the current vector distribution. Several other genes detected in the selective sweep analysis have been implicated in the host response to apicomplexan protozoa (which includes *Theileria* species), which lends credence to the hypothesis that the ancient co-evolution and selection pressure exerted by *T. parva* in *S. caffer* may have played a role in shaping the patterns of diversity in relevant regions of the current *S. caffer* genome. The long relationship between *T. parva* and *S. caffer* is reflected in the limited pathology caused by infection of *T. parva* in *S. caffer*, which is in stark contrast to the severe and often fatal disease caused by *T. parva* infection in other hosts such as domestic cattle [18, 62]. The latter have only co-existed with *T. parva* for 5,000-10,000 years [63]. This finding may provide a route to identifying genes and pathways important in controlling disease during infections by *Theileria* species, that can, for example, be translated to mitigating the effect of these pathogens upon cattle or Asian buffalo owned by resource-poor farmers.

## Conclusion

For the first time we have analysed genome-level data from all extant recognised African buffalo subspecies, covering the majority of the remaining geographical distribution of the species. Our findings demonstrate that the African buffalo is composed of three main lineages, that further subdivide based on geographical location. While current subspecies nomenclature is likely to still have utility in terms of Management or Conservation Units, more samples and data, particularly from *S. c. nanus*, *S. c. brachyceros* and *S. c. aequinoctialis*, would help resolve the status of taxonomic units across the population range of African buffalo. The data also demonstrated that genetic connectivity between populations has historically been constrained by geographical barriers that have shaped the modern population structure (particularly the Congo basin), and that human influence has been for ∼10,000 years and remains a main pressure on effective population size and population fragmentation. While most populations do not show signs of inbreeding, particular populations do, and this has implications for conservation and management of the species. Finally, through analyses of selective sweeps, we identified infectious diseases as a likely substantial contributor to historical selection, and hypothesise that protozoan pathogens for which the buffalo has been primary host for millennia may be responsible for driving some of this selection.

## Materials & Methods

### Sample collection

DNA samples were obtained through (1) active sampling of animals for this project; this was done in collaboration with the Kenya Wildlife Service at the Ol Pejeta Conservancy, Kenya, or (2) secondary use of DNA samples previously collected; this included samples previously collected and published from Tanzania [14], Uganda [64], and Mozambique, Botswana, Zimbabwe, South Africa, Niger, Burkina Faso, Gabon Central African Republic and Chad [5, 6, 8]. For sample collection in Kenya, buffalo were darted and sedated by qualified veterinary personnel from KWS, and 10 ml blood collected into Paxgene Blood DNA tubes from peripheral venous sampling. DNA was extracted from the Paxgene Blood DNA tubes using the Paxgene Blood DNA kit (Qiagen) according to the manufacturer’s instructions. Tissue pieces (OPB4) were snap frozen in liquid nitrogen in the field. Tissue pieces were homogenised using mortar and pestle over liquid nitrogen. The powder was resuspended in Trireagent (Sigma-Aldrich) and RNA was isolated using the RNeasy kit (Qiagen) according to the manufacturer’s instructions.

Relevant research approvals were obtained in all instances; for the active sampling within this study, approval was obtained from the Kenya Wildlife Services (permit number KWS/BRM/5001). For secondary use of DNA samples previously collected, relevant permits are Tanzania Wildlife Research Institute and Tanzania Commission for Science and Technology (permit number 2021-262-NA-2021-066) [14] and Uganda Wildlife Authority (permit number COD/96/05) [64], or details are provided in [5, 6, 8].

### Genome sequencing

For the reference genome, a buffalo sample from Ol Pejeta in Kenya (OPB4) was sequenced using a combination of Illumina HiSeq (Dovetail Genomics & Edinburgh Genomics) and Pacific BioSciences approaches (Dovetail Genomics & Edinburgh Genomics) to a final sequencing coverage of 75× (Illumina) and 60× (PacBio). The same sample was also sequenced using Illumina Hi-C (Dovetail Genomics) in order to facilitate scaffolding. For the population samples, approximately 2.5 µg of total DNA from 196 animals sampled across Africa (Kenya, Uganda, Tanzania, Mozambique, Botswana, Zimbabwe, South Africa, Niger, Burkina Faso, Gabon, Central African Republic and Chad; Table 1, Supplementary Table 2) was subjected to whole-genome sequencing by Illumina HiSeq; this was performed at a coverage of 30× for 50 samples from Tanzania, with the remaining samples being sequenced at 15×.

### Genome assembly

A primary assembly of the single molecule PacBio sequencing from OPB4 (mean read lengths > 10Kb) was generated using FALCON and consisted of 7,269 contigs and an N50 of 1.9 Mb. This primary assembly was scaffolded using the Hi-C libraries and the HiRise software by Dovetail. The resulting scaffold-level assembly was further improved via gap filling and polishing steps performed with PBJelly [27] and Pilon [28] respectively, as described below. Gap filling: 7,085 gaps (both inter– and intra-scaffolds) were identified in the scaffold-level assembly. A total of 78 inter-scaffold gaps were partially filled (i.e. extended on one side) using PBJelly, with 476,665 bases added in total, while none of the identified gaps were fully closed. This observation confirmed the high quality of the primary assembly achieved from PacBio reads including a post-processing step using Arrow (part of the GenomicConsensus package from PacBio). Polishing: An additional 75× Illumina short read sequencing (101bp paired-end reads) of DNA from the same individual used to build the reference genome assembly (OPB4), was used to polish the *de novo* scaffold-level reference genome assembly. Polishing allows the correction of artefacts due to sequencing errors in assemblies, using the pile up of short reads that are associated with low sequencing error (∼1%). This process was performed multiple times and improvement upon quality metrics (i.e. reduced numbers of ambiguous bases, corrected SNPs, resolved small indels, closed gaps) were assessed after each round of Pilon (see Supplementary Data 1). The rate of improvements reached a plateau between the third (P3) and the fourth (P4) rounds of Pilon, and therefore the resulting P4 polished assembly was considered optimal and used for downstream analysis. Given the reference genome should not contain any homozygote alternate variant calls relative to the short read data from the same sample, we compared how the number of these changed following polishing. The Illumina short reads, sequenced from the same animal as that used to generate the reference genome assembly (OPB4), were mapped with bwa-mem (BWA v0.7.17) against the polished genome assemblies (P2 to P4). The percentages of mapped reads were extremely high (>99%) and comparable across the P2, P3 and P4 assemblies.

### Assembly statistics

To directly compare the quality of the genome assembly at each step during the assembly process, and to highlight improvements, QUAST (v 5.0.2) [65] was used to produce genome assembly metrics for each iteration of the genome assembly, pre and post gap filling with PBJelly, and for each successive round of polishing with Pilon (Supplementary Data 1). QUAST further compares a given genome assembly to a reference genome, and for this the genome assembly for the water buffalo *Bubalus bubalis* (GCF_003121395.1) [29] was provided, to produce genome alignment metrics and details of suspected misassemblies (Supplementary Data 1). A custom Python script (https://raw.githubusercontent.com/evotools/CattleGraphGenomePaper/master/Assembly/ABS.py) was used to calculate scaffold metrics, N, L, NG, LG and GC content for a given proportion of the scaffold-level P4 genome assembly, in 5% increments (5-100, Supplementary Data 1). The scaffold-level P4 genome assembly contains a total of 3,351 scaffolds, of which 1,381 scaffolds are greater than 10 kb. Quality values (QV) representative of the single-base accuracy were computed using Merqury (v1.1)[66]with the K-mer counts generated by Meryl (v1.2; https://github.com/marbl/meryl). For downstream analysis we selected 1,381 contigs with a length of 10 kb or greater, representing 99.68% (2.653 Gb) of the total length of the assembled genome. This subset of contigs were used for downstream analyses.

### Detection of novel genomic sequences

Following completion of the assembly, we identified the novel sequences in the genome in comparison with other ruminant species. We selected a set of nine genome assemblies for five species, and calculated the distances among them using mash v2.2 [67], using a K-mer size of 32. We used the following genome assemblies to generate the alignment graph: *Syncerus caffer* (accession number GCA_902825105.1), *Bubalus bubalis* Mediterranean (GCF_003121395.1)[29], *Capra hircus* San Clemente (GCF_001704415.1)[34], *Bos grunniens* (GCA_005887515.2)[30], *Bos taurus indicus* Brahman (GCF_003369695.1), *Bos taurus taurus* Angus (GCA_003369685.2)[31], *Bos taurus taurus* Hereford (GCF_002263795.1)[33], *Bos taurus taurus* N’Dama (GCA_905123515) and *Bos taurus indicus* Ankole (GCA_905123885)[32]. We then generated a phylogenetic tree using the neighbour-joining algorithm included in the neighbour software from Phylip (v3.698)[68] which was used to create the following guide tree for CACTUS [69]:

((angus:0.00187,hereford:0.00115)Anc1:0.0004,(ankole:0.00317,((yak:0.00671,((abuffalo:0. 01228,wbuffalo:0.0095)Anc6:0.00438,goat:0.04443)Anc5:0.01195)Anc4:0.00254,brahman:0.00256)Anc3:0.00023)Anc2:0.0004,ndama:0.00195)Anc0;

The HAL archive of multiple whole genome alignments (mWGA) was generated using the software CACTUS [69], and then converted to PackedGraph format using the hal2vg software (v.2.1)[70] with the African buffalo genome as reference. We then used the nf-GraphSeq workflow (https://github.com/evotools/CattleGraphGenomePaper/tree/master/detectSequences/nf-GraphSeq) described in Talenti et al. [32] based on libbdsg [71] to identify the nodes (i.e. the fragment of genome) that are found exclusively in the backbone of the graph (i.e. African buffalo genome), excluding all intervals overlapping a gap. We combined all interval regions less than 5bp apart using BEDTools (v.2.30.0) [72]. We then annotated the regions by length (short if < 10bp, intermediate if < 60bp and large if > 60bp), position (labelled telomeric if <10Kb from the end of a scaffold larger than 5Mb, flanking a gap if <1Kb from an N-mer), type of sequence (novel if > 95% of the bases in the region are not found in any other genome, or haplotype if < 95% of the bases were found only in the African Buffalo) and proportion of masked bases. We filtered out regions if they 1) were not classified as long, 2) contained less than 50% novel bases and 3) were not telomeric or were not flanking a gap.

To validate that these regions corresponded to buffalo sequence, and did not derive, for example, from contamination, 46 of the population WGS samples were randomly selected and their coverage examined at these regions, with the assumption that if these regions corresponded to contamination in our reference sample, they would not have aligned reads from multiple buffalo samples. Mean read depth was calculated for each of the 74,659 novel regions within the reference genome, for the 46 population samples, using Mosdepth (v0.3.4) [73]. The distribution of average coverage values across the population samples, for each novel region, is shown in Supplementary Figure 1. There are only 1494 novel regions with a mean read depth < 1 and 419 regions with no reads mapped across these 46 samples, suggesting that these putative African buffalo-specific regions do not derive from an artefact such as contamination.

We characterized the content of the novel regions by 1) performing a motif analysis using HOMER (v4.11.1)[74], and 2) by detecting the novel features. To identify these features, we used the annotation generated by Ensembl and available in the rapid release database (http://www.ensembl.info/2020/06/25/ensembl-rapid-release/; accession GCA_902825105.1). We identified all gene features overlapping a novel sequence using bedtools intersect (v2.30.0) [72], and identified only these fully overlapping a novel region still with bedtools intersect with the –f 1.0 option (100% of overlap between the feature and the novel region).

Once we identified these fully new gene features, we extracted the GO term and KEGG pathways present in the annotation itself in embl format. To do so, we first converted the file in GenBank format, and then extracted for each gene the transcript IDs, protein IDs and biological terms. For these terms, we performed an enrichment analysis in R using a binomial test with the genes not in novel regions as background.

### Reference genome annotation

Genome annotation was undertaken at EMBL-EBI by Ensembl, primarily using RNA-seq and full-length isoform sequencing (Iso-Seq) data generated from the animal for which the genome was assembled. A TruSeq stranded total RNA-seq library with one round of Ribo-Zero Gold kit (Illumina) was prepared from one pooled library consisting of RNA samples from eight tissues (heart, prescapular and inguinal lymph nodes, testis, liver, kidney, lung and spleen) collected from the animal for which the genome was assembled. RNA-seq was performed at Edinburgh Genomics on an S2 lane of an Illumina NovaSeq 6000 platform generating 100bp paired-end reads. Iso-Seq was performed at the Centre for Genomic Research at the Univerity of Liverpool, using RNA samples from six different tissues (prescapular lymph node, testis, liver, kidney, lung and spleen) collected from the same animal. Full-length cDNA from total RNA was generated using TeloPrime full-length cDNA amplification kit (v2) from Lexogen. A total of six barcoded TeloPrime libraries from six RNA samples were multiplexed. Iso-seq was performed on the resulting multiplexed library using six PacBio Sequel SMRT cells. The RNA-seq data were aligned to the reference genome using STAR [75]. For loci where the structures derived from the transcriptomic data appeared to be fragmented or absent, gap-filling using cross-species protein data was carried out. For more information on the annotation process see Supplementary Information 1.

### Detection of variants in WGS samples across Africa

For all 196 WGS samples from *S. caffer* across Africa (raw data is available at ENA via accession numbers PRJEB59220 and ERP144275), reads were mapped with bwa-mem (BWA v0.7.17) against the reference genome generated as above. The GATK (v4.0.11.0) pipeline, following the best practices as outlined at https://gatk.broadinstitute.org/hc/en-us/articles/360036194592-Getting-started-with-GATK4 was used with HaplotypeCaller to identify variants (SNPs and Indels). The GATK best practice includes a Variant Quality Score Recalibration (VQSR) step that compares all variant calls to those in a high quality set to identify and flag potential false positives. Unlike in well-characterised species no gold-standard set of variants is available for the African buffalo. We therefore used a consensus set of 6,806,905 variants called from the Illumina data generated for the same sample as the reference genome using three software tools (GATK, Arrow and Longshot [76]). Although we do not expect this set to be free of false variant calls, we expect it to be enriched for true positives and was therefore used in VQSR. Three VQSR tranches, 99, 99.9 and 100 (each representing the proportion of gold-standard variants that are retained at each quality threshold), were assessed. The variant set resulting from the 99.9 tranche was selected for downstream analyses with a Ti/Tv ratio of 2.07 and >120M variants. The variant set was further filtered for GQ (Phred-scaled Probability that the call is incorrect) values less than 30 and site missingness of 0.9 (at least 90% of the samples contain data at this site). PLINK (v1.90) was used to calculate sample missingness, the proportion of variant sites missing from each sample, and vcftools (v0.1.13) to calculate the relatedness of all individuals. For downstream analyses, individual samples with a missingness greater than 0.15, additionally individuals that were closer than fourth degree relatedness (relatedness value 0.0625), were also removed, resulting in a variant dataset covering 163 individual animals. We checked for any mapping biases due to use of an East African reference genome, by randomly sampling three animals per country and comparing how read mapping rates differed by longitude (Supplementary Figure 7). No obvious mapping bias was observed among the West African samples when mapping to the reference genome obtained from an East African sample.

### Genomic diversity analyses

The VCF file for the set of unrelated samples was first filtered through bcftools (v1.9; https://samtools.github.io/bcftools/) to keep only unrelated individuals according to the KING method implemented in vcftools [77, 78]. A cutoff of 0.0625 was applied to exclude 3^rd^ degree relatives or closer. Furthermore only biallelic SNPs in large contigs (>10Kb) were retained. Variants were further filtered using plink (v1.90b4) [79] to restrict to those with a minor allele frequency >0.05. This dataset was then used to carry out analyses of migration events and effective population size. ADMIXTURE and the identity-by-state phylogenetic tree can benefit from having an even sample size for the different populations/samples deriving from the same location that were tested [38]. Therefore, for these analyses we identified a representative subsample for the populations with more than 15 animals. Sample size reduction was carried out using the BITE R package [80] to select a representative set of individuals for each population. The reduction process was performed on each population separately. For each group we selected the variants with very high call rate (99%) and highly polymorphic (--maf 0.3). The reduction step in BITE was performed considering only individuals with 95% call rate and up to 10K markers to compute the kinship matrix (options n.trials = 100000, ibs.marker=10000, n.k=2, ibs.thr = 0.95, id.cr=0.95). Principal component analysis (PCA) was performed post reduction in sample numbers using plink v1.90b4. Admixture analysis was performed using ADMIXBoots (https://github.com/RenzoTale88/ADMIXBoots), a Nextflow workflow that performs bootstrapped admixture (v1.3.0) [81], defining a consensus of the different K at different iterations using CLUMPP [82] and generating plots in R. The workflow was run pruning for variants in linkage (plink –-indep-pairwise 5000 100 0.3), testing every K between 2 and 15, and with 100 bootstraps of 100,000 markers each. A consensus of the different bootstraps was called using CLUMPP in LargeKGreedy mode. Bar charts for each consensus K, boxplots for the distribution of the CV errors and line plots of the H’ scores of each K were generated from the pipeline automatically. Bootstrapped identity-by-state (IBS) phylogenetic tree was calculated using the nf-PhyloTree workflow (https://github.com/RenzoTale88/nf-PhyloTree). This workflow uses plink to generate a matrix of identity-by-state distances across individuals. These workflows then use a series of custom scripts to generate the individual phylogenetic tree, call the consensus tree and generate the input compliant for GraphLan [83]. In our case, we ran the workflow allowing for pruning variants in high linkage disequilibrium using plink (--indep-pairwise 5000 100 0.3), then generated 100-bootstrapped IBS-based distances using plink (--distance 1-ibs square flat-missing), allowing repeated variants in each bootstrap and sampling a number of SNPs equalling the number of pruned variants. A consensus tree was generated using a custom python script, converted to phyloXML, annotated for colour and consistency nodes and finally plotted with GraphLan [83]. For the isolation by distance analysis, pairwise F_ST_ values between populations were calculated using vcftools, and the Haversine formula was used to calculate the distances between the centre points of population sampling sites.

### Estimated Effective Migration Surfaces (EEMS)

The EEMS package developed by Petkova et al. [84] was used (https://github.com/dipetkov/eems) to estimate effective migration surfaces. The runeems_snps program was used to visualise spatial population structure in the African buffalo populations and to identify the geographic barriers to migration preventing gene flow across these populations. The runeems_snps program requires the following data as input files: (1) a matrix of average pairwise genetic dissimilarities, (2) sample coordinates, and (3) a list of habitat coordinates, here covering the natural distribution of African buffalo populations on the African continent, and listed as a sequence of vertices organised as a closed polygon. For the input files for EEMS analysis, a matrix of average pairwise genetic dissimilarities was generated from the pruned set of SNP data, using the bed2diffs_v1 program within the EEMS package. The locations of all African buffalo animals, from which DNA samples were collected for WGS and variant detection, were inputted as longitude and latitude coordinates, indicating either specific sampling locations or the centre of specific geographical regions (*e.g.* national parks) when no other information was available. The list of habitat coordinates was generated based on the known past and present natural distribution of the four subspecies of African buffalo populations (as described in [2]) and using the https://www.latlong.net/ website to identify the latitude and longitude geocoding of point locations on the African continent. EEMS analysis was run using the runeems_snps program within the EEMS package based on the African buffalo pruned SNP data. Parameters used to run EEMS analysis were set as follows: nIndiv = 163; nSites = 6000; nDemes = 400; diploid = true; numMCMCIter = 4000000; numBurnIter = 1000000; numThinIter = 9999. Description for all parameters used are defined in the EEMS instruction manual (v.0.0.0.9000). Results of EEMS analysis were plotted using the rEEMSplot package in R to generate contour plots of effective migration and effective diversity surfaces from EEMS outputs. Additionally, posterior probability trace plots (pilogl) were used to check the MCMC sampler had successfully converged using four million MCMC iterations. The effective migration and diversity surfaces plots also include the addition of lakes and rivers depicted in blue based on data extracted from the Natural Earth website (https://www.naturalearthdata.com/download/50m/physical/).

### Estimating effective population sizes and selective sweeps

To calculate the XP-EHH scores the African buffalo genotype data was first phased using Beagle 5.1 [85]. A recombination rate of 1cM/Mb was assumed and XP-EHH scores calculated between each pair of populations using hapbin [86]. Peaks were called as previously described [42]. Briefly XP-EHH scores were smoothed by averaging across 1000 SNP windows and putative selective sweep regions were those with an absolute XP-EHH >4, with the start and end coordinates defined where the XP-EHH scores fell back below two. The locations of XP-EHH peaks in the water buffalo and cattle genomes were obtained from Dutta et al. [42] and the peaks for all three species mapped to the orthologous regions of the water buffalo genome.

The effective population sizes over time of the three largest African buffalo populations were calculated using Relate v1.1.6 [40] using the same phased haplotypes from Beagle. An estimated generation time of 11 years for the African buffalo was used in this analysis [87]. Previously calculated estimated effective population sizes for human African populations were obtained from Speidel et al [40]. The P_R_ statistic was also calculated using Relate [40] and the same Beagle haplotype files using an estimated mutation rate of 1.25×10^-8^. Variants with a P less than 5×10^-8^ were retained. The circular Manhattan plot was created using the CMplot R package [88]. The water buffalo genes were lifted over to the African buffalo genome to identify which genes fell under putative selective sweep peaks.

### Data availability statement

All raw data generated in this project is available through the following routes: for the buffalo reference genome, PacBio, Illumina and Hi-C data is available via Genbank (GCA_902825105.1) and ENA (PRJEB3658), and the assembly, annotation and associated flat files can be accessed through https://rapid.ensembl.org/Syncerus_caffer_GCA_902825105.1; transcriptomic data were deposited to ENA (https://www.ebi.ac.uk/ena/browser/view/GCA_902825105.1) with accession numbers PRJEB36587 and PRJEB36588 for RNA-seq and Iso-Seq, respectively, and population genome data is available through ENA via accession number PRJEB59220.

## Acknowledgements

This research was funded in part by the Bill & Melinda Gates Foundation (USA) and with UK aid from the UK Foreign, Commonwealth and Development Office (Grant Agreement OPP1127286) under the auspices of the Centre for Tropical Livestock Genetics and Health (CTLGH), established jointly by the University of Edinburgh (UK), Scotland’s Rural College (SRUC, UK), and the International Livestock Research Institute (UK). The findings and conclusions contained within are those of the authors and do not necessarily reflect positions or policies of the Bill & Melinda Gates Foundation nor the UK Government. The work was also supported by grants BBS/OS/GC/000012C, BBS/E/D/20002172 and BBS/E/D/20002174 from the Biotechnology and Biological Sciences Research Council (BBSRC, UK). This research was conducted as part of the Consultative Group on International Agricultural Research (CGIAR) Program on Livestock (Kenya). International Livestock Research Institute (ILRI), Kenya, is supported by contributors to the CGIAR Trust Fund. CGIAR is a global research partnership for a food-secure future. Its science is carried out by 15 Research Centres in close collaboration with hundreds of partners across the globe (www.cgiar.org). Work carried out at EMBL, EBI was supported by the Wellcome Trust (WT222155/Z/20/Z) and BBSRC (BB/S020152/1).

## Supplementary Data 1 Genome assembly statistics

**Supplementary Figure 1.**
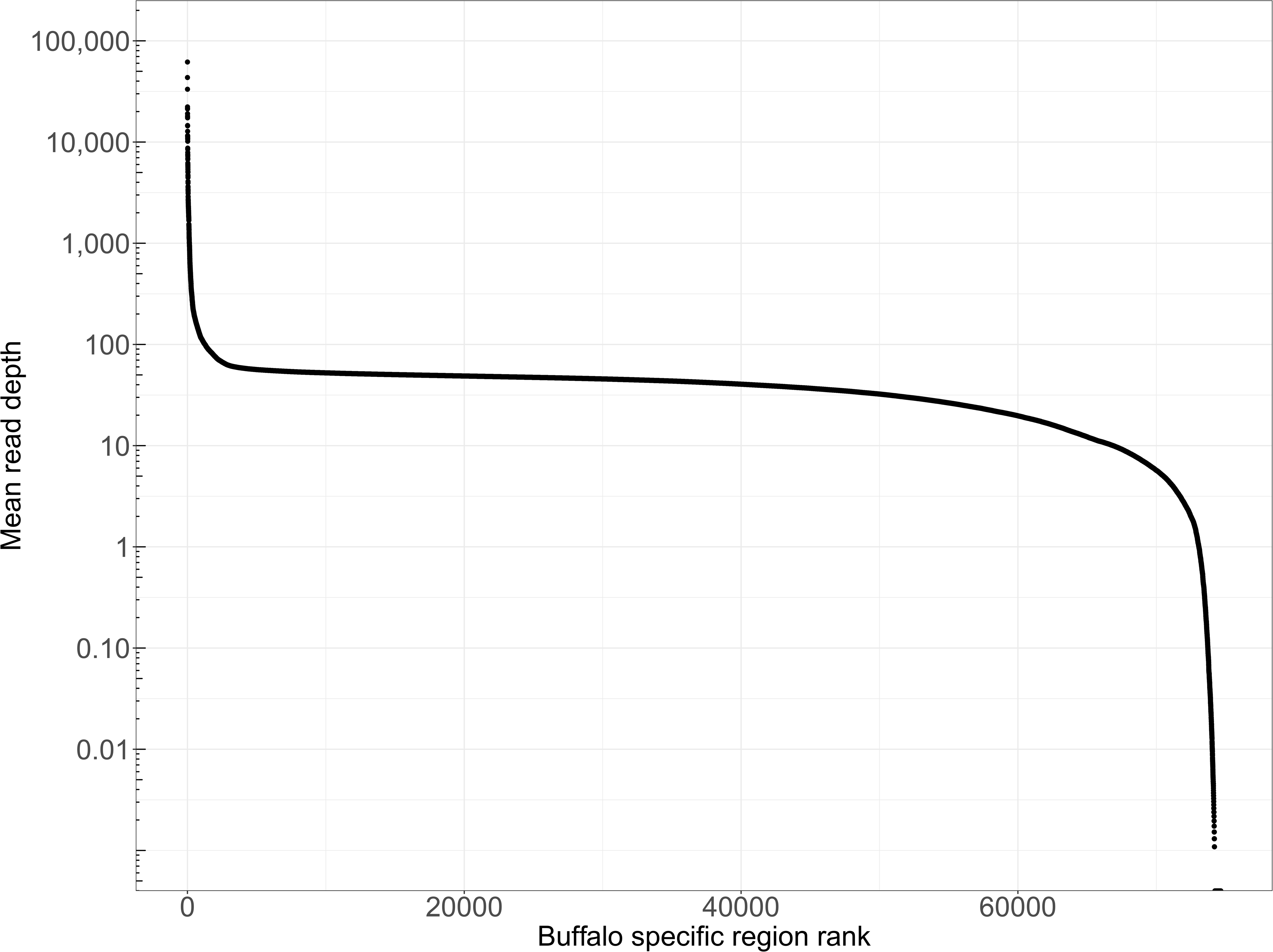
Mean read depth across the putative African buffalo specific regions. Mean read depth was calculated for each of the 74,659 novel regions within the reference genome, for 46 of the population samples, using Mosdepth (v0.3.4) [73]. The distribution of average coverage values across the population samples, for each novel region, is shown. There are only 1494 novel regions with a mean read depth < 1 and 419 regions with no reads mapped across these 46 samples.

Supplementary Table 1. Genes identified in the buffalo-specific sequence, with Ensembl transcript, gene and protein identifiers, and GO terms where relevant. Note that the list is greater than the 583 identifed genes, as some genes appear in the list more than once due to having different transcripts.

Supplementary Table 2. Details of buffalo samples for which genome sequences were generated and included in this study, including sample identification, subspecies, country of origin, region of origin, whether sequences were retained in analysis following filtering steps (0.0625 relatedness, 0.2 missingness), the population group the sample was assigned to, and a latitude/longitude of a central point in the respective sampling area.

**Supplementary Figure 2.**
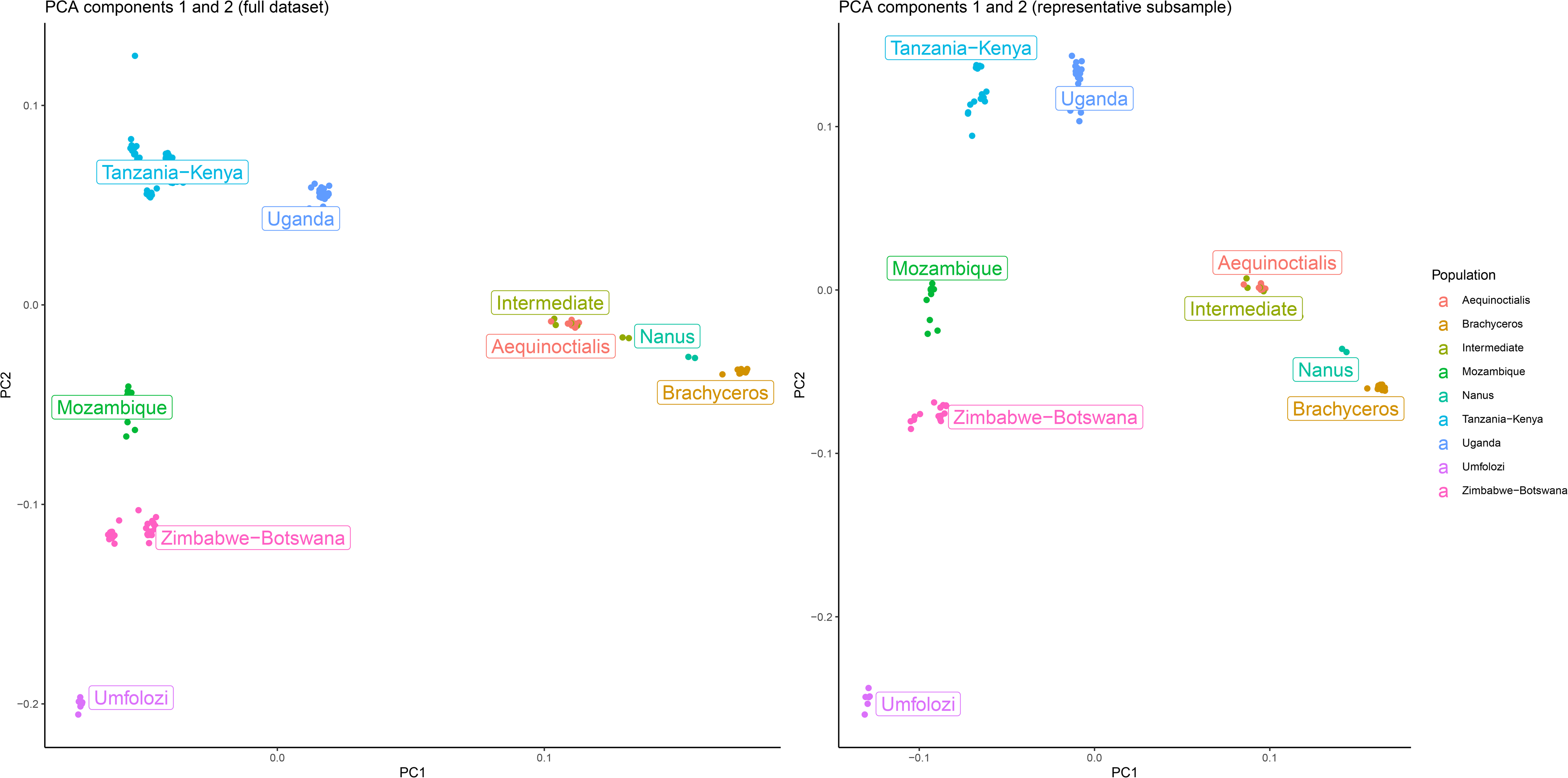
Principal Component analysis pre-& post-reduction (i.e. following sample removal post filtering steps; 0.0625 relatedness, 0.2 missingness), with data for components 1 and 2 illustrated, samples are coloured by population grouping.

**Supplementary Figure 3.**
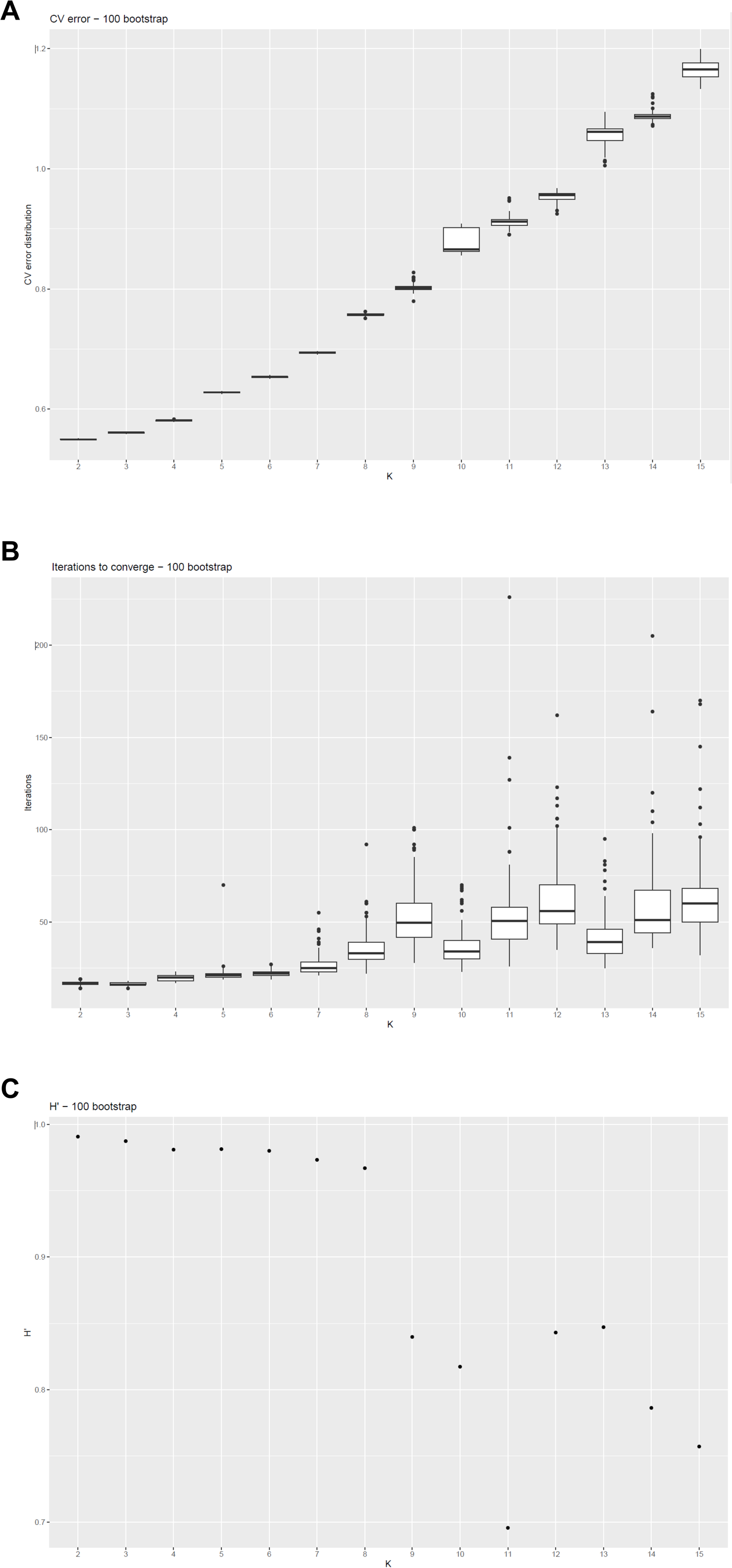
Admixture evaluation metrics ((A) cross-validation error, (B) number of iterations to converge and (C) H’) at different values of K calculated using 100 bootstraps of 100,000 variants each.

**Supplementary Figure 4.**
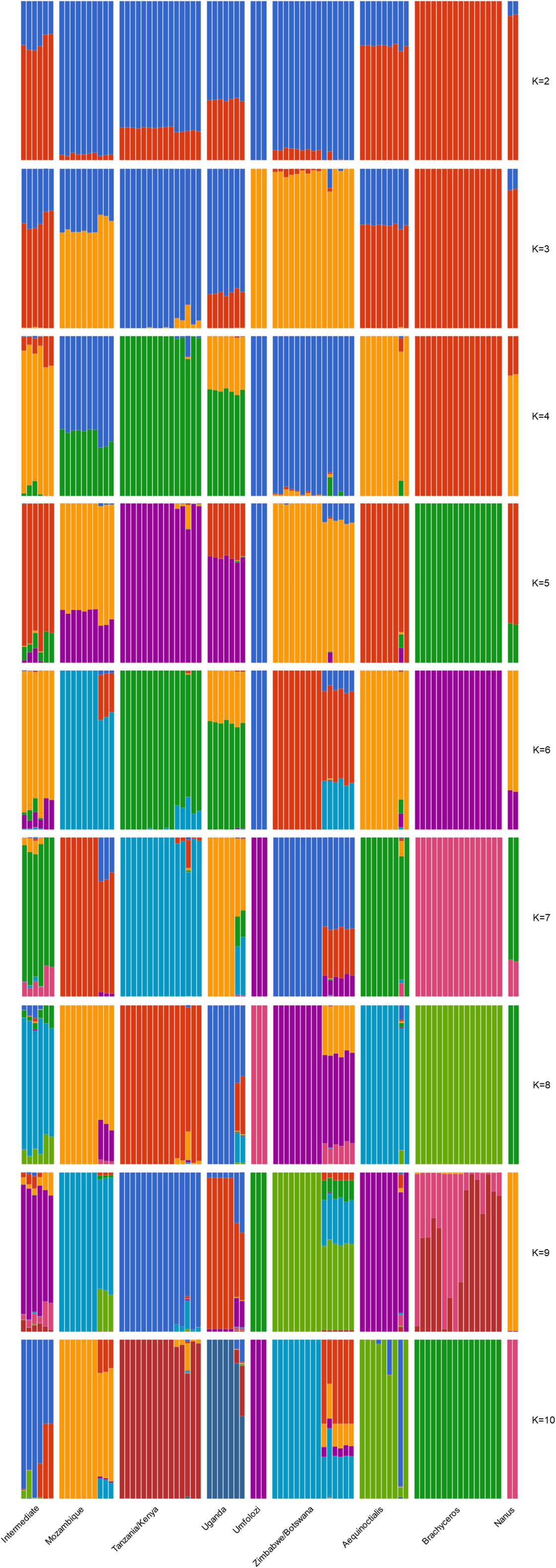
Admixture analysis for K = 2-10.

Supplementary Table 3. Pairwise F_ST_ values for the nine population groupings (*S. c brachyceros*, *S. c. nanus*, *S. c. aequinoctialis*, intermediate (putative hybrids between *S. c. nanus*, *S. c. aequinoctialis*), *S. c. caffer* Uganda, *S. c. caffe*r Kenya/Tanzania, *S. c. caffer* Mozambique, *S. c. caffer* Zimbabwe/Botswana and *S. c. caffer* South Africa), and geographic distance as measured to centred latitude/longitude measurement for each grouping.

Supplementary Table 4. Details of genes identified to be under selection in the African buffalo, whether the gene has been previously identified to be in a selection peak in either the cow or water buffalo, and whether the gene is related to immune response function. Genes are grouped by (a) detected in both XPEHH and PR analyses of African buffalo (dark green), (b) detected in either XPEHH or PR analyses of African buffalo, and in both metrics for water buffalo or cow analyses (medium green), (c) detected in either XPEHH or PR analyses of African buffalo, and in one of the metrics for water buffalo or cow analyses (light green), or (d) none of the above (no colour).

**Supplementary Figure 5.**
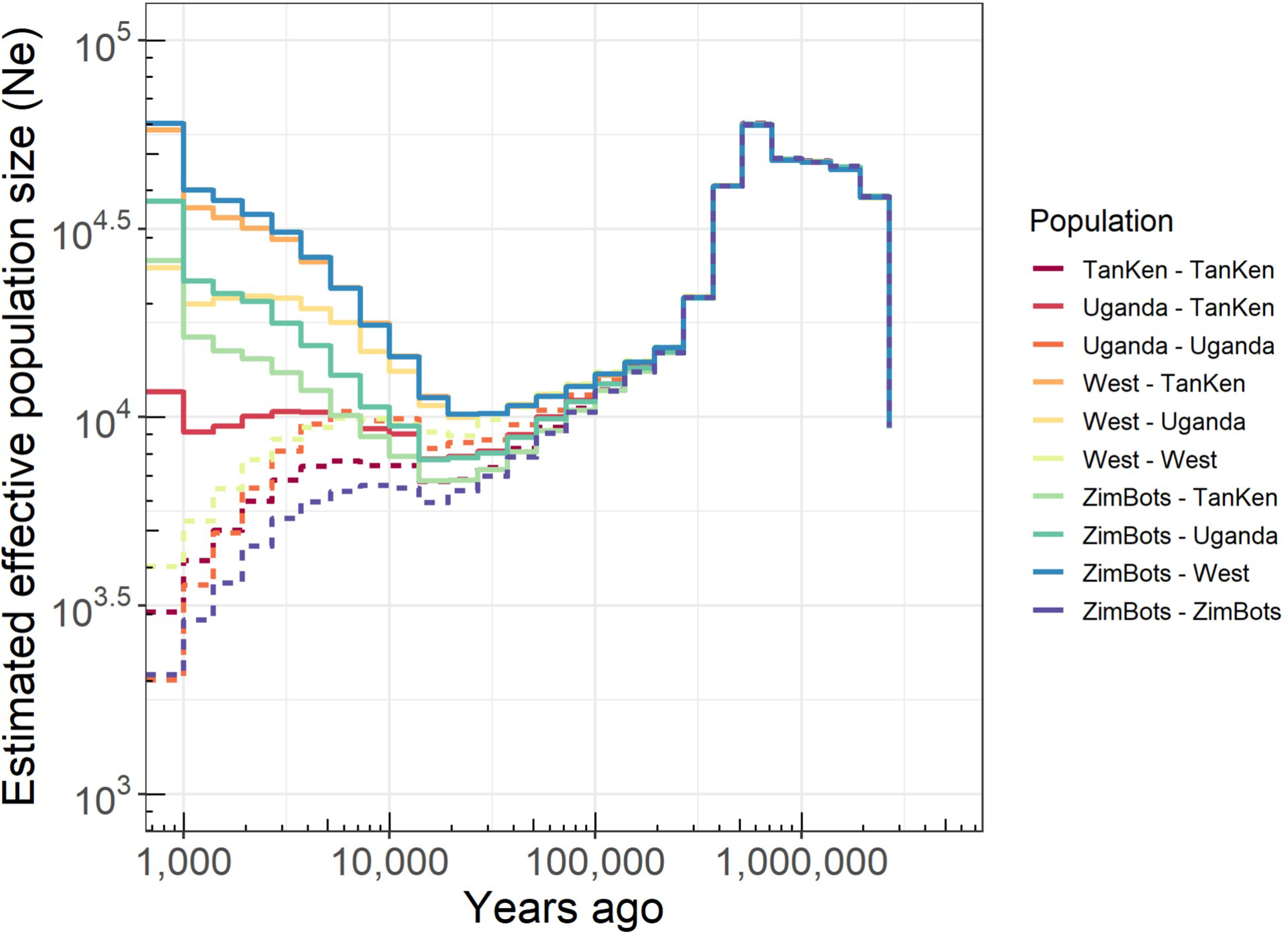
Relate-inferred inverse coalescence rates (effective population sizes) for each of the larger sub-groups to themselves (dashed lines) and each other (solid lines). For this comparison, due to the smaller sample sizes, all West African animals were collated into one group.

**Supplementary Figure 6.**
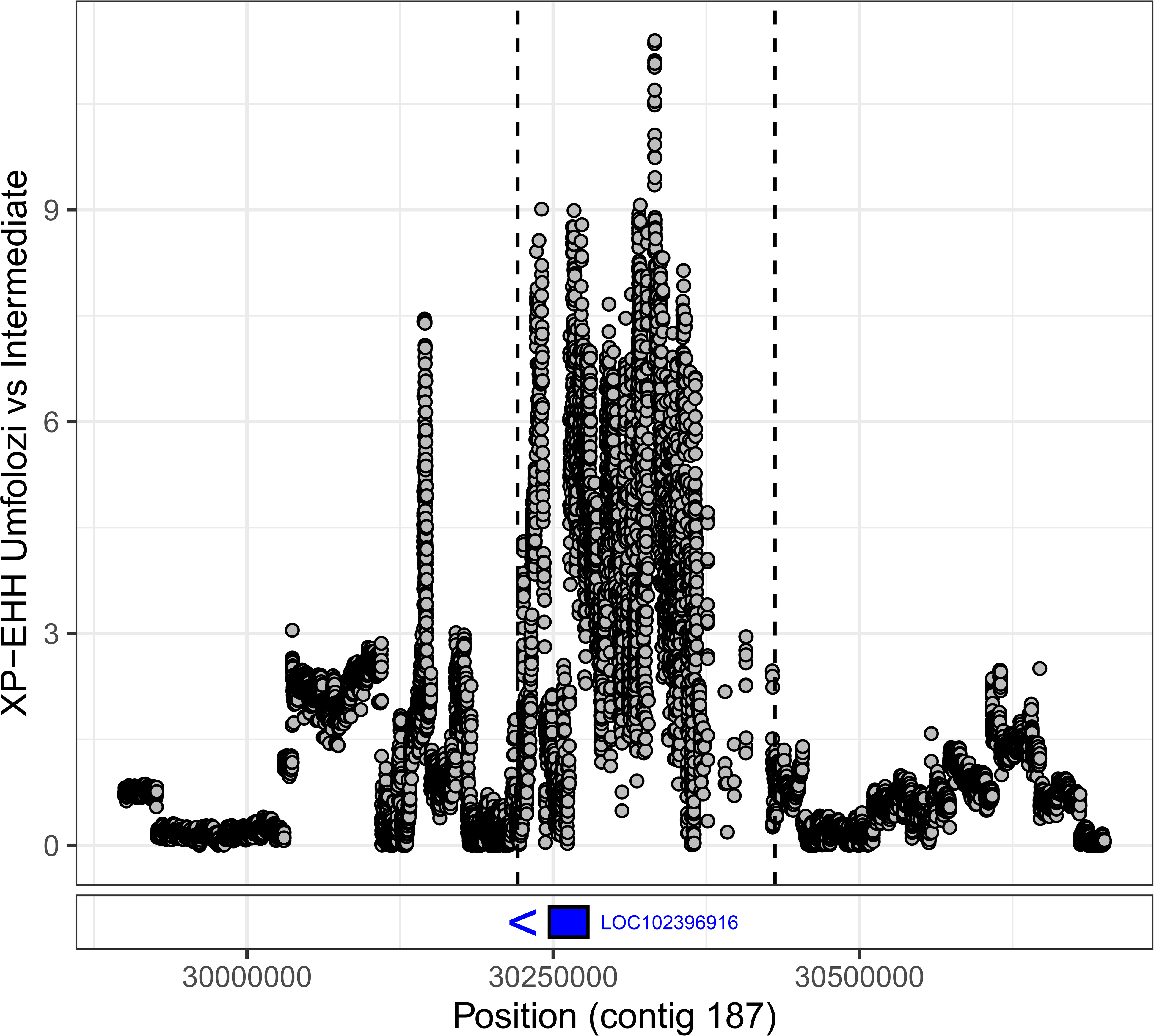
The absolute XP-EHH scores at the LOC102396916 locus on contig 187. The boundaries of the called peak are indicated by dashed vertical lines. The location and direction of LOC102396916 is shown in blue below.

Supplementary Information 1 A methodological summary of the genome annotation process undertaken at ENSEMBL.

**Supplementary Figure 7.**
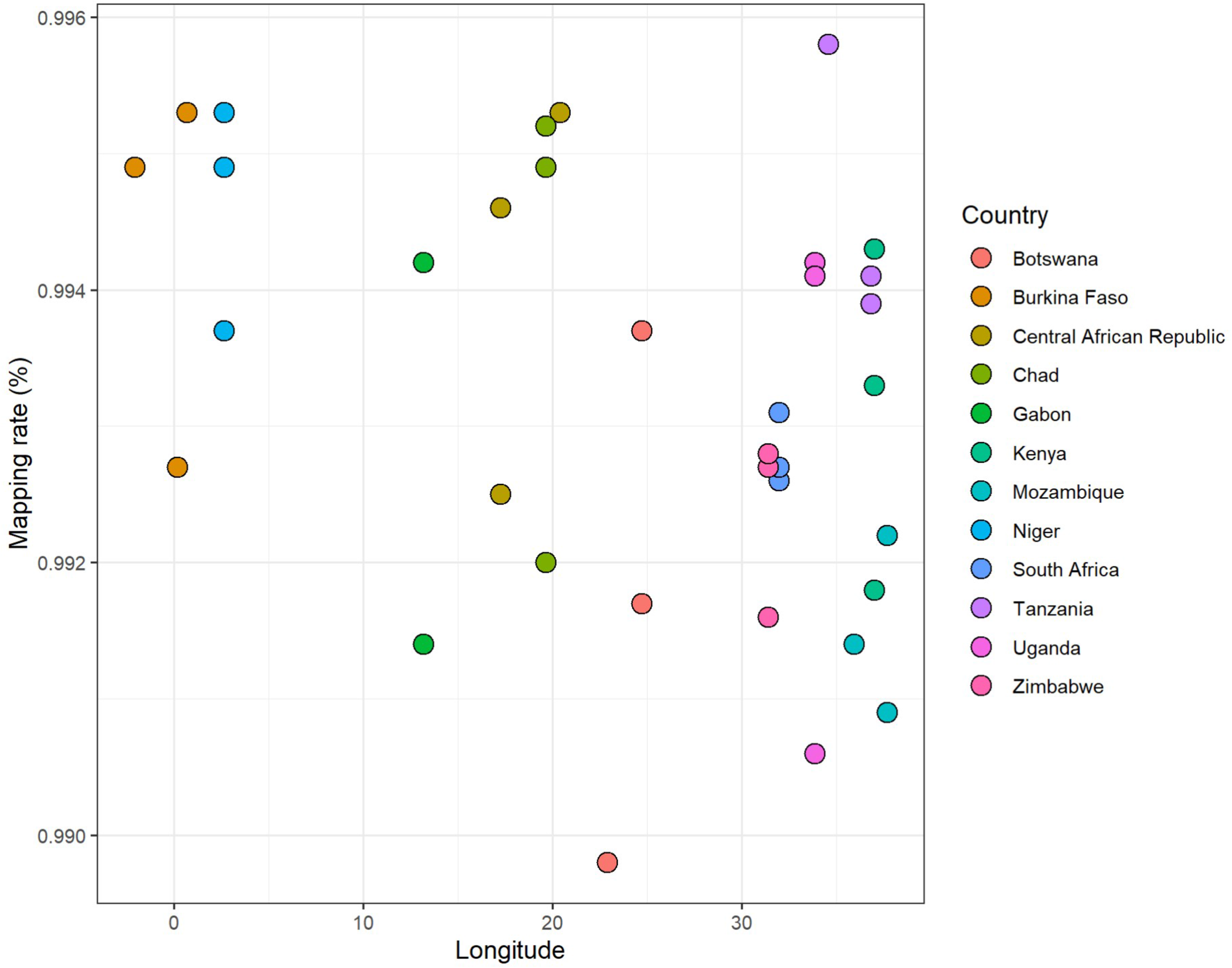
Mapping rate by longitude of three randomly selected samples per country. No obvious mapping bias was observed among the West African samples when mapping to the reference genome obtained from an East African sample.

