## Supplementary Data 1 for "Continent-wide genomic analysis of the African buffalo (*Syncerus caffer*)"

Supplementary Data 1, Table 1. Assembly statistics at each round of polishing.

| Assembly | Unfilled | Filled | pilon1 | pilon2 | pilon3 | pilon4 |
| --- | --- | --- | --- | --- | --- | --- |
| # sequences (>= 0 bp) | 3,351 | 3,351 | 3,351 | 3,351 | 3,351 | 3,351 |
| # sequences (>= 1000 bp) | 3,041 | 3,042 | 3,043 | 3,043 | 3,043 | 3,043 |
| # sequences (>= 5000 bp) | 2,136 | 2,157 | 2,156 | 2,155 | 2,154 | 2,154 |
| # sequences (>= 10000 bp) | 1,352 | 1,378 | 1,379 | 1,380 | 1,380 | 1,381 |
| # sequences (>= 25000 bp) | 320 | 325 | 325 | 325 | 325 | 325 |
| # sequences (>= 50000 bp) | 208 | 208 | 209 | 209 | 211 | 211 |
| Total length | 2,651,354,003 | 2,651,819,263 | 2,652,421,646 | 2,652,507,212 | 2,652,733,160 | 2,652,945,513 |
| GC (%) | 41.76 | 41.76 | 41.75 | 41.75 | 41.75 | 41.75 |
| Scaffold N50 | 69,170,419 | 69,170,419 | 69,160,108 | 69,159,231 | 69,159,740 | 69,160,875 |
| N75 | 31,468,510 | 31,468,510 | 31,463,761 | 31,459,774 | 31,459,818 | 31,459,638 |
| L50 | 13 | 13 | 13 | 13 | 13 | 13 |
| L75 | 27 | 27 | 27 | 27 | 27 | 27 |
| # N's per 100 kbp | 149 | 149 | 123 | 116 | 113 | 112 |

Supplementary Data 1, Table 2. QUAST Assembly metrics. Contig length distribution, and N50 metrics for the genome assembly prior and post PBJelly gap filling and after each successive round of  polishing with Pilon.

| Assembly | Unfilled | Filled | Pilon1 | Pilon2 | Pilon3 | Pilon4 |
| --- | --- | --- | --- | --- | --- | --- |
| # contigs (>= 0 bp) | 3,351 | 3,351 | 3,351 | 3,351 | 3,351 | 3,351 |
| # contigs (>= 1000 bp) | 3,041 | 3,042 | 3,043 | 3,043 | 3,043 | 3,043 |
| # contigs (>= 5000 bp) | 2,136 | 2,157 | 2,156 | 2,155 | 2,154 | 2,154 |
| # contigs (>= 10000 bp) | 1,352 | 1,378 | 1,379 | 1,380 | 1,380 | 1,381 |
| # contigs (>= 25000 bp) | 320 | 325 | 325 | 325 | 325 | 325 |
| # contigs (>= 50000 bp) | 208 | 208 | 209 | 209 | 211 | 211 |
| Largest contig | 193,316,640 | 193,316,640 | 193,289,354 | 193,281,852 | 193,285,335 | 193,288,002 |
| Total length | 2,651,354,003 | 2,651,819,263 | 2,652,421,646 | 2,652,507,212 | 2,652,733,160 | 2,652,945,513 |
| N50 | 69,170,419 | 69,170,419 | 69,160,108 | 69,159,231 | 69,159,740 | 69,160,875 |

Supplementary Data 1, Table 3. QUAST Assembly metrics compared against *Bubalus bubalis* reference genome. Genome length, alignment and assembly metrics for the genome assembly prior and post PBJelly gap filling and after each successive round of polishing with Pilon.

| Assembly | Unfilled | Filled | Pilon1 | Pilon2 | Pilon3 | Pilon4 |
| --- | --- | --- | --- | --- | --- | --- |
| Reference length | 2.65 Gb | 2.65 Gb | 2.65 Gb | 2.65 Gb | 2.65 Gb | 2.65 Gb |
| GC (%) | 41.76 | 41.76 | 41.75 | 41.75 | 41.75 | 41.75 |
| Reference GC (%) | 41.81 | 41.81 | 41.81 | 41.81 | 41.81 | 41.81 |
| # misassemblies | 10,283 | 10,289 | 55,205 | 55,190 | 55,180 | 55,219 |
| # misassembled contigs | 594 | 592 | 1,024 | 1,021 | 1,027 | 1,028 |
| # unaligned contigs | 357 + 1377 part | 358 + 1413 part | 265 + 823 part | 265 + 834 part | 266 + 829 part | 261 + 833 part |
| Unaligned length | 445,613,097 | 446,253,486 | 73,191,862 | 73,130,730 | 73,194,004 | 73,190,626 |
| Genome fraction (%) | 82 | 82 | 94 | 94 | 94 | 94 |
| # N's per 100 kbp | 149 | 149 | 123 | 116 | 113 | 112 |
| # mismatches per 100 kbp | 4,324 | 4,325 | 2,104 | 2,104 | 2,104 | 2,104 |
| # indels per 100 kbp | 202 | 202 | 258 | 256 | 256 | 255 |
| Largest alignment | 6,016,346 | 6,016,343 | 1,669,457 | 1,669,468 | 1,669,432 | 1,669,453 |
| Total aligned length | 2.2 Gb | 2.2 Gb | 2.571 Gb | 2.571 Gb | 2.572 Gb | 2.572 Gb |
| NA50 | 674,686 | 674,686 | 128,221 | 128,284 | 128,283 | 128,332 |
| NGA50 | 673,117 | 673,117 | 127,916 | 128,119 | 128,113 | 128,156 |

Supplementary Data 1, Table 4. Genome Assembly Scaffold Metrics. N, L, NG, LG and GC content for a given proportion of the genome assembly, x.

| x | Nx | Lx | NGx | LGx | GC% |
| --- | --- | --- | --- | --- | --- |
| 5 | 193,288,002 | 0 | 193,288,002 | 0 | 40.17 |
| 10 | 143,084,846 | 1 | 143,084,846 | 1 | 40.99 |
| 15 | 136,240,748 | 2 | 136,240,748 | 2 | 40.87 |
| 20 | 111,633,540 | 3 | 111,633,540 | 3 | 41.03 |
| 25 | 104,800,038 | 4 | 104,800,038 | 4 | 40.87 |
| 30 | 94,608,040 | 6 | 94,608,040 | 6 | 40.6 |
| 35 | 89,027,109 | 7 | 89,027,109 | 7 | 40.6 |
| 40 | 73,376,136 | 9 | 73,376,136 | 9 | 40.72 |
| 45 | 71,999,969 | 10 | 71,869,821 | 11 | 40.73 |
| 50 | 69,160,875 | 12 | 66,444,531 | 13 | 40.82 |
| 55 | 63,542,860 | 14 | 63,539,877 | 15 | 41.17 |
| 60 | 56,889,046 | 16 | 55,993,668 | 17 | 41.38 |
| 65 | 47,354,412 | 19 | 43,700,853 | 20 | 41.33 |
| 70 | 39,306,693 | 22 | 38,815,046 | 23 | 41.33 |
| 75 | 31,459,638 | 26 | 26,790,382 | 27 | 41.46 |
| 80 | 25,550,784 | 31 | 23,490,963 | 32 | 41.62 |
| 85 | 19,164,195 | 37 | 15,575,171 | 39 | 41.65 |
| 90 | 8,757,406 | 46 | 6,900,340 | 52 | 41.65 |
| 95 | 3,798,400 | 68 | 1,747,249 | 84 | 41.73 |
| 100 | 2 | 3,351 | 2 | 3351 | 41.7 |
