## Supplementary Information 1 for "Continent-wide genomic analysis of the African buffalo (*Syncerus caffer*)"

**Ensembl Genome Annotation**

Annotation of the assembly was created via the Ensembl gene annotation system (PMID: 27337980). A set of potential transcripts was generated using multiple techniques: primarily through alignment of transcriptomic data sets, and also through gap filling with protein-to-genome alignments of a sub-set of mammalian proteins from UniProt (PMID: PMC6323992). The UniProt mammalian proteins had experimental evidence for existence at the protein or transcript level (protein existence level 1 and 2). Additionally, a whole genome alignment was generated between the genome and the GRCh38 human reference genome using LastZ and the resulting alignment was used to map the coding regions of human genes from the GENCODE reference set.

At each locus, low quality transcript models were removed, and the data were collapsed and consolidated into a final gene model plus its associated non-redundant transcript set. When collapsing the data, priority was given to models derived from transcriptomic data. For each putative transcript, the coverage of the longest open reading frame was assessed in relation to known vertebrate proteins, to help differentiate between true isoforms and fragments. In loci where the transcriptomic data were fragmented or missing, homology data was used to gap fill if a more complete cross-species alignment was available, with preference given to longer transcripts that had strong intron support from the short-read data.

Gene models were classified, based on the alignment quality of their supporting evidence, into three main types: protein-coding, pseudogene, and long non-coding RNA. Models with hits to know proteins, and few structural abnormalities (i.e., they had canonical splice sites, introns passing a minimum size threshold, low level of repeat coverage) were classified as protein-coding. Models with hits to known protein, but having multiple issues in their underlying structure, were classified as pseudogenes. Single-exon models with a corresponding multi-exon copy elsewhere in the genome were classified as processed pseudogenes.

If a model failed to meet the criteria of any of the previously described categories, did not overlap a protein-coding gene, and had been constructed from transcriptomic data then it was considered as a potential lncRNA. Potential lncRNAs were filtered to remove transcripts that did not have at least two valid splice sites or cover 1000bp (to remove transcriptional noise).

A separate pipeline was run to annotation small non-coding genes. miRNAs were annotated via a BLAST (PMID: 2231712) of miRbase (PMID:30423142) against the genome, before passing the results in to RNAfold (PMID: 18424795). Poor quality and repeat-ridden alignments were discarded. Other types of small non-coding genes were annotated by scanning Rfam (PMID: 29112718) against the genome and passing the results into Infernal (PMID: 24008419).

The annotation for the African buffalo is available via Ensembl Rapid Release: <https://rapid.ensembl.org/Syncerus_caffer_GCA_902825105.1/Info/Index>
